# Feature processing for single-trial semantic decoding from EEG: a factorial comparison with pre-registered replication

**DOI:** 10.64898/2026.11.20.768234

**Authors:** Laurence Yang, Milad Lankarany

## Abstract

Single-trial decoding of semantic processing from EEG is used in cognitive neuroscience, brain–computer interfaces and EEG-to-text research. Because single trials are noisy, the choice of features can determine how much information a decoder uses. Yet these fields use different features that have not been compared systematically: event-related potential studies decode the voltage time course (the waveform), EEG-to-text studies mostly the power of oscillations in frequency bands. We crossed four feature-processing choices—signal representation (waveform or band power), time window, normalization and channel set—into 640 configurations and decoded whether a stimulus fitted its semantic context in three tasks: visual word pairs, objects in visual scenes, and words in spoken sentences (40 participants per task). Eleven pre-specified comparisons were tested jointly. The signal representation had by far the largest effect on accuracy (mean generalized *η*^2^ = 0.165, an order of magnitude above any other effect). Averaged over tasks and other choices, balanced accuracy was 56.6% with the waveform and 50.6% with theta-band power (chance, 50%), and adding band power to the waveform lowered accuracy by 1.8 percentage points, in every task. Both effects replicated, and were larger, in a pre-registered study of 283 new participants from three sites. Neither a finer time–frequency decomposition including the delta band (2–4 Hz) nor a different way of combining power with the waveform changed this conclusion, whereas the time window, normalization and channel set mattered little. For semantic decoding from single EEG trials, the waveform carries the information, and band power does not add to it.

## 1 Introduction

The N400 component of the event-related potential (ERP) is the most widely used electrophysiological index of semantic processing: words, pictures and objects that fit poorly into their context evoke a larger negativity that peaks around 400 ms after their onset and, for words, is largest over centro-parietal sites (Kutas and Hillyard, 1980; Kutas and Federmeier, 2011). Increasingly, such effects are analysed not only in trial averages but with multivariate decoding, which asks whether the experimental condition can be read out from the spatiotemporal pattern of the EEG (Grootswagers et al., 2017; Carrasco et al., 2024). Semantic decoding has been used to detect lexico-semantic processing in individual children (Petit et al., 2020), to trace the time course of negation effects (He et al., 2024), as a step towards N400-based brain–computer interfaces (Dijkstra et al., 2020; Wimmer et al., 2024) and, in a different community, as the input to models that translate EEG into text (Wang and Ji, 2022). Because the signal-to-noise ratio of single trials is low, the way the EEG is turned into features before classification can determine how much of the available information a decoder is able to use.

The fields that decode semantic information from EEG have, however, settled on very different features. ERP-oriented studies typically use the broadband voltage, averaged in short time bins and concatenated across channels (Blankertz et al., 2011). EEG-to-text studies have mostly inherited the features distributed with the ZuCo corpus: the power in eight frequency bands, aggregated over the fixations on each word (Hollenstein et al., 2018; Wang and Ji, 2022). Whether such models exploit the EEG at all has recently been questioned (Jo et al., 2024), but the more basic question of which signal representation carries single-trial semantic information has not been addressed.

The answer is not obvious. Semantic violations are accompanied not only by the N400 but also by changes in theta and gamma power (Hald et al., 2006), so band power could carry information that complements the waveform. In other domains the two representations can carry dissociable information: sustained potentials carry information about both the location and, independently of it, the orientation of an item held in working memory, whereas alpha-band power carries information about the location but little or none about the orientation (Bae and Luck, 2018). For object categories, a systematic comparison of EEG features found wavelet coefficients about as informative as the raw voltage samples, but the Hilbert amplitude less so (Karimi-Rouzbahani et al., 2021). Whether band power adds to the broadband waveform in semantic decoding is therefore an empirical question.

Decoding studies of the three kinds of semantic context examined here have also each used a single representation, almost always the voltage. For visual word pairs, the N400 effect has been decoded from the voltage at each time point, in single trials and in averages of several trials (Trammel et al., 2023), and the ERP CORE N400 data have been decoded in the same format in the methodological studies described below (Kessler et al., 2025; Zhang et al., 2024). For objects in scenes, the consistency of an object with its scene was decoded from the voltage at each time point (Draschkow et al., 2018b), the same responses have been analysed with representational similarity analysis of voltage patterns (Krugliak et al., 2024), and object categories in consistent and inconsistent scenes have been decoded from short voltage segments (Chen et al., 2022). For sentences, congruency has been decoded from the voltage at each time point in spoken sentences, in part of the data analysed here (Heikel et al., 2018), and in written sentences (He et al., 2024). Among the decoding studies we found, only one used band power: it concatenated spectral features with window-averaged amplitudes to detect semantic violations in spoken sentences and did not evaluate the two separately (Tanaka et al., 2019). In these paradigms, power has otherwise been examined in condition averages (Hald et al., 2006), so whether it adds to the voltage in single trials has not been tested.

Methodological work on ERP decoding does not yet answer it. On the ERP CORE dataset (Kappenman et al., 2021), which includes an N400 experiment, a multiverse analysis crossed eight preprocessing steps, from re-referencing and filtering to artifact correction, and showed that they affect decoding performance both individually and in interaction (Kessler et al., 2025). Other studies on the same data evaluated one choice at a time: artifact correction and rejection (Zhang and Luck, 2025), spatial principal component analysis (Zhang et al., 2024), classifier regularization and cross-validation (Zhang et al., 2026a), sampling rate (Zhang et al., 2026b) and spatial normalization (Qin et al., 2026); electrode density has been examined in other paradigms (Hou et al., 2026). For the N400, classifiers have been compared on the ERP CORE data and a second word-priming experiment (Trammel et al., 2023) and in an augmented-reality task (Wimmer et al., 2024), without varying the features. These studies varied mainly how the EEG is cleaned or classified; where they changed the features, they examined a single step, such as spatial principal component analysis, spatial normalization or the number of electrodes, and none of them varied the signal representation. The other feature-processing choices that every decoding analysis has to make—the time window, the normalization of the features and the set of channels—have thus been examined at most one at a time and not tested for interactions. Moreover, conclusions drawn from one dataset need not hold for other semantic tasks. To our knowledge, no study has compared these choices factorially for semantic decoding, or tested at the single-trial level whether band power carries semantic information beyond the waveform.

Here we compared feature processing for single-trial semantic decoding factorially and across tasks (Fig. 1). We crossed four dimensions—signal representation (the broadband waveform, amplitude envelopes in four frequency bands and their combinations), time window, normalization and channel set—into 640 configurations, and decoded semantic congruency within participants in three tasks that differ in sensory modality and in the kind of semantic context: visual word pairs from ERP CORE, and objects in visual scenes and words in spoken sentences from a second dataset (Draschkow et al., 2018a,b; Heikel et al., 2018). Eleven pre-specified contrasts were tested jointly across the three tasks, and each effect was classified as general, consistent in direction but varying in size, task-dependent, or without a reliable general effect. The two most robust conclusions were then tested in a pre-registered replication on an independent sample of 283 participants recorded at three sites, and a pre-specified time–frequency analysis tested whether they reflected a power representation that was too coarse or unfairly combined with the waveform.

**Figure 1:**
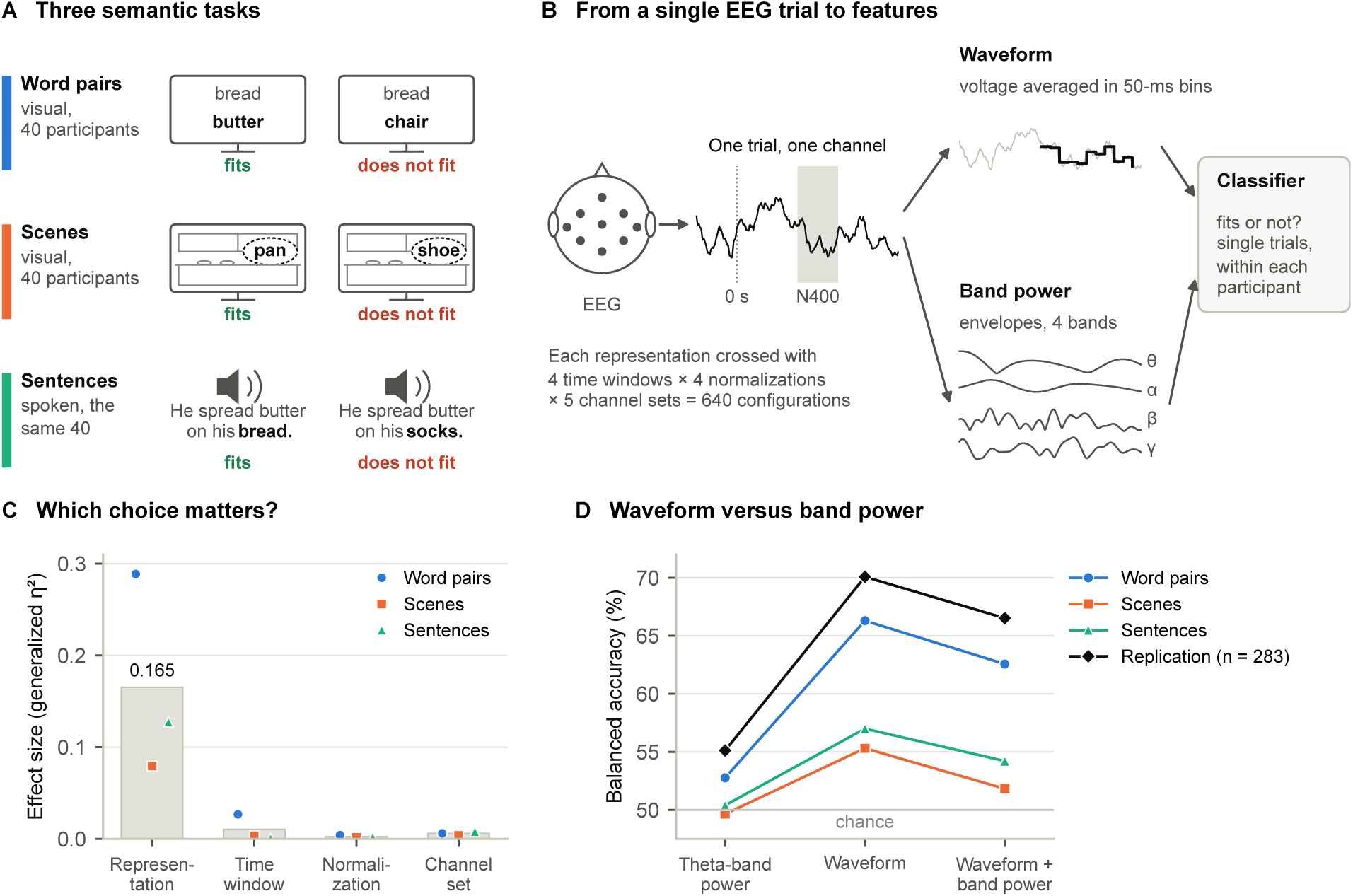
Overview of the study. (A) Semantic congruency was decoded in three tasks: visual word pairs (ERP CORE), objects in scenes and spoken sentences; the scene and sentence tasks were completed by the same 40 participants. The examples are schematic; the sentences were in German. (B) Each EEG trial was represented by its waveform (the voltage, averaged in 50-ms bins) or by the power in frequency bands (the amplitude envelopes in the theta, alpha, beta and low-gamma bands), alone or in combination, and these representations were crossed with four time windows, four normalizations and five channel sets (640 configurations). A classifier was trained within each participant to tell from single trials whether the stimulus fitted its context. The trial shown is simulated. (C) Effect of each feature-processing choice on accuracy: generalized *η*^2^ of the four main effects (bars, mean of the three tasks; symbols, single tasks). (D) Mean balanced accuracy with theta-band power (B2), the waveform (B1) and the waveform combined with the four band envelopes (B8) at the baseline settings (200–800 ms, per-feature standardization, all channels; logistic regression), in the three tasks and in the pre-registered replication. Confidence intervals and tests are given in Figs. 5 and 6.

From the prior work and common practice, we expected the waveform to outperform single-band envelopes, and treated as open whether band power adds information to the waveform. We further expected a late time window to add little to the classical N400 window, per-trial normalization not to improve on per-feature standardization, multivariate noise normalization to perform at least as well as per-feature standardization (Guggenmos et al., 2018), and all channels to perform at least as well as the centro-parietal N400 region, which in turn should outperform frontal and posterior regions.

## 2 Materials and methods

### 2.1 Datasets

We analysed three semantic-congruency tasks drawn from two openly available datasets, and used a third, independent dataset for the replication (Table 1). All tasks contrast two classes (congruent vs. incongruent; chance level 50%), and all decoding was performed within participants.

**Table 1:**
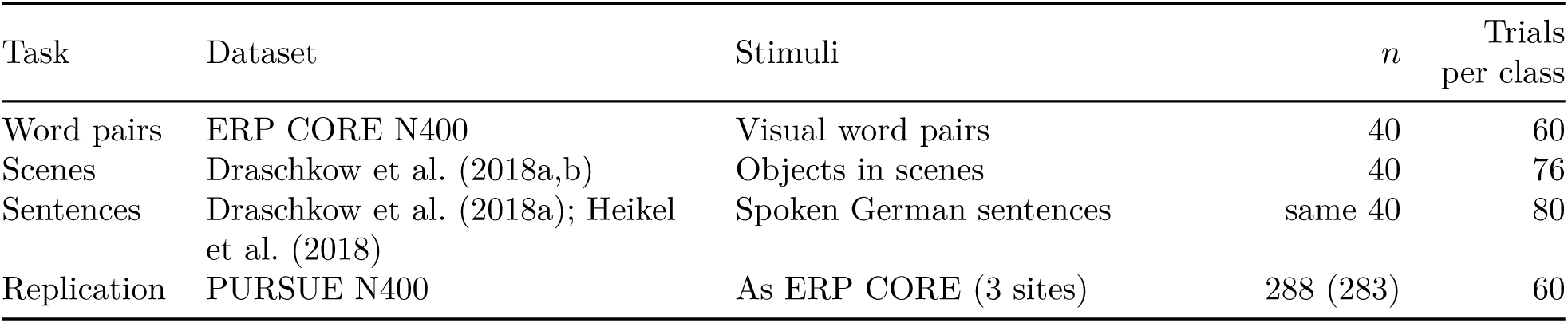
Datasets. *n*, number of participants (for the replication, the number analysed in parentheses); trials per class are those recorded per participant.

#### ERP CORE N400 (word pairs)

The N400 experiment of ERP CORE (Kappenman et al., 2021; Luck, 2020) comprises 40 participants recorded with a BioSemi ActiveTwo system at 1024 Hz from 30 scalp electrodes and three ocular electrodes. On each trial a red prime word (200 ms) was followed after 900–1100 ms by a green target word (200 ms) that was either related or unrelated to the prime; participants indicated the relation by pressing one of two buttons as quickly as possible. Each participant completed 120 trials (60 per class), and each target word appeared once in each class. Event times were shifted by the measured display delay of 26 ms.

#### Visual scenes and spoken sentences

The dataset of Draschkow et al. (2018a,b) comprises 40 participants who completed both tasks in one session; it is independent of ERP CORE. EEG was recorded at 1000 Hz with Brain Products actiCHamp or BrainAmp amplifiers; about half of the participants were recorded with 62 channels referenced online to the linked mastoids, and the others with 64 channels, which additionally contained the two mastoid electrodes. In the scene task, an object appeared in a scene with which it was either consistent or inconsistent (76 trials per class); trials were locked to the object markers. In the sentence task, participants listened to German sentences whose critical word was either congruent or incongruent with the sentence context (80 trials per class, half with high and half with low cloze probability); Heikel et al. (2018) analysed the sentences with high cloze probability. Critical-word onsets were computed with the rule in the authors’ code (condition marker + 255 ms audio delay + the word onset listed for that sentence) and agreed with the authors’ epochs to within 6 ms (median 1 ms) in all 40 participants, with identical class labels. Neither task required a speeded judgement of congruency. All 28 non-reference electrodes used by our ERP CORE channel sets were present.

#### PURSUE N400 (replication)

The PURSUE N400 word-processing dataset (Couperus et al., 2026) contains 288 participants recorded at three undergraduate institutions (Mount Holyoke College, University of Richmond, Claremont McKenna College) with the ERP CORE word-pair paradigm and event codes. EEG was recorded with an actiCHamp system at 500 Hz from 29 scalp electrodes and three ocular channels. The PURSUE participants do not overlap with those of the other datasets.

#### Ethics

The present work is a secondary analysis of de-identified, openly available data collected in earlier studies. The collection of all three datasets was approved by the responsible ethics committee or institutional review boards, and all participants gave informed consent, as reported by the original authors (Kappenman et al., 2021; Draschkow et al., 2018b; Heikel et al., 2018; Couperus et al., 2021).

### 2.2 Preprocessing

Preprocessing followed one fixed path, the example path of Kessler et al. (2025), which approximates the original ERP CORE preprocessing, so that the comparison concerned feature processing alone. Data were downsampled to 256 Hz; bipolar horizontal and vertical EOG channels were computed; the data were provisionally referenced to Cz; ocular and muscle artifacts were removed with two independent component analyses (Picard; 20 components; components identified with the MNE-Python EOG and muscle detectors) (Ablin et al., 2018a,b; Gramfort et al., 2013); the data were high-pass filtered at 0.1 Hz (no low-pass filter) and re-referenced to the mean of P9 and P10. Epochs from *−*0.4 to 0.8 s around the target were baseline-corrected (*−*0.2 to 0 s), and bad channels within epochs were interpolated with autoreject (Jas et al., 2017) without rejecting epochs. We made two implementation-level changes: the rare epochs in which every channel was flagged, and which autoreject therefore sets to zero, were removed, and the band envelopes (Section 2.3) were interpolated on the analytic signal rather than on the envelope, so that envelopes could not become negative. All target trials entered the analysis regardless of response accuracy.

For the scene and sentence tasks, the path was adapted where the recordings required it. The data were already referenced to the linked mastoids (for 64-channel recordings, after converting to the mean of M1 and M2), which we kept as the final reference; the provisional Cz reference was therefore skipped. The recorded ocular channels SO1 and SO2 served as EOG, and no display delay was applied to the scene task because none was reported. The saved epochs and all features were restricted to the 28 electrodes that are also used for ERP CORE (independent component analysis and interpolation used all channels). For the sentence task, the muscle independent component analysis was omitted, and instead of interpolation, epochs were rejected if their peak-to-peak amplitude exceeded a global threshold estimated per participant with autoreject’s cross-validated procedure, after removing flat epochs (median channel standard deviation below 1 µV). The restriction to 28 electrodes and the two sentence-task settings were adjusted after a three-participant pilot check (Supplementary Section S2), and the flat-epoch criterion was added after one recording was found to be partly flat (Supplementary Section S7). Re-running the full analysis with the originally specified settings left all joint categories of the planned contrasts (Section 2.6) unchanged for logistic regression; for shrinkage linear discriminant analysis (LDA; Section 2.5), only the classification of P2, which was not robust in the main analysis, changed (Section 3.7).

For the replication, the ERP CORE path was applied unchanged except for the final reference, which was the mean of TP9 and TP10 (the authors’ offline reference; P9 and P10 were not recorded). Features used the 27 of the 28 ERP CORE electrodes that were recorded (FCz was missing).

### 2.3 Band-limited amplitude envelopes

To avoid edge effects in short epochs, band-limited signals were computed on the continuous preprocessed data. Each recording was band-pass filtered (FIR) in the theta (4–8 Hz), alpha (8– 13 Hz), beta (13–30 Hz) and low-gamma (30–45 Hz) bands, and the analytic signal was obtained with the Hilbert transform. Epochs from *−*0.5 s to the end of the waveform epoch were extracted for the same trials, the channels flagged by autoreject in a trial were interpolated on the real and imaginary parts, and the modulus gave the amplitude envelope. Envelopes were log-transformed and baseline-corrected per trial and channel by subtracting the mean log amplitude between *−*500 and *−*200 ms; this window lies before the temporal smearing of the theta-band filter reaches the stimulus. Following common usage, we refer to these log amplitude features, and to the log wavelet amplitudes of Section 2.8, as band power; log power is twice the log amplitude, so the two differ only by a constant factor.

### 2.4 Feature-processing dimensions

The design is summarized in Fig. 2. Each trial is represented as an array of components *×* channels *×* time, where a component is either the waveform or a band envelope. Features were computed in a fixed order: selection of the components (representation), log transform and baseline correction of envelopes, selection of the time window and averaging in 50-ms bins, selection or combination of channels, normalization, and flattening into a feature vector. The four dimensions and their levels are listed in Table 2; full crossing gives 8 *×* 4 *×* 4 *×* 5 = 640 configurations, with B1-W1-N2-C1 (waveform, 200–800 ms, per-feature standardization, all 28 electrodes) as the baseline, which follows the conventional ERP decoding approach of averaging the broadband voltage in short time bins on all channels (Blankertz et al., 2011).

**Figure 2:**
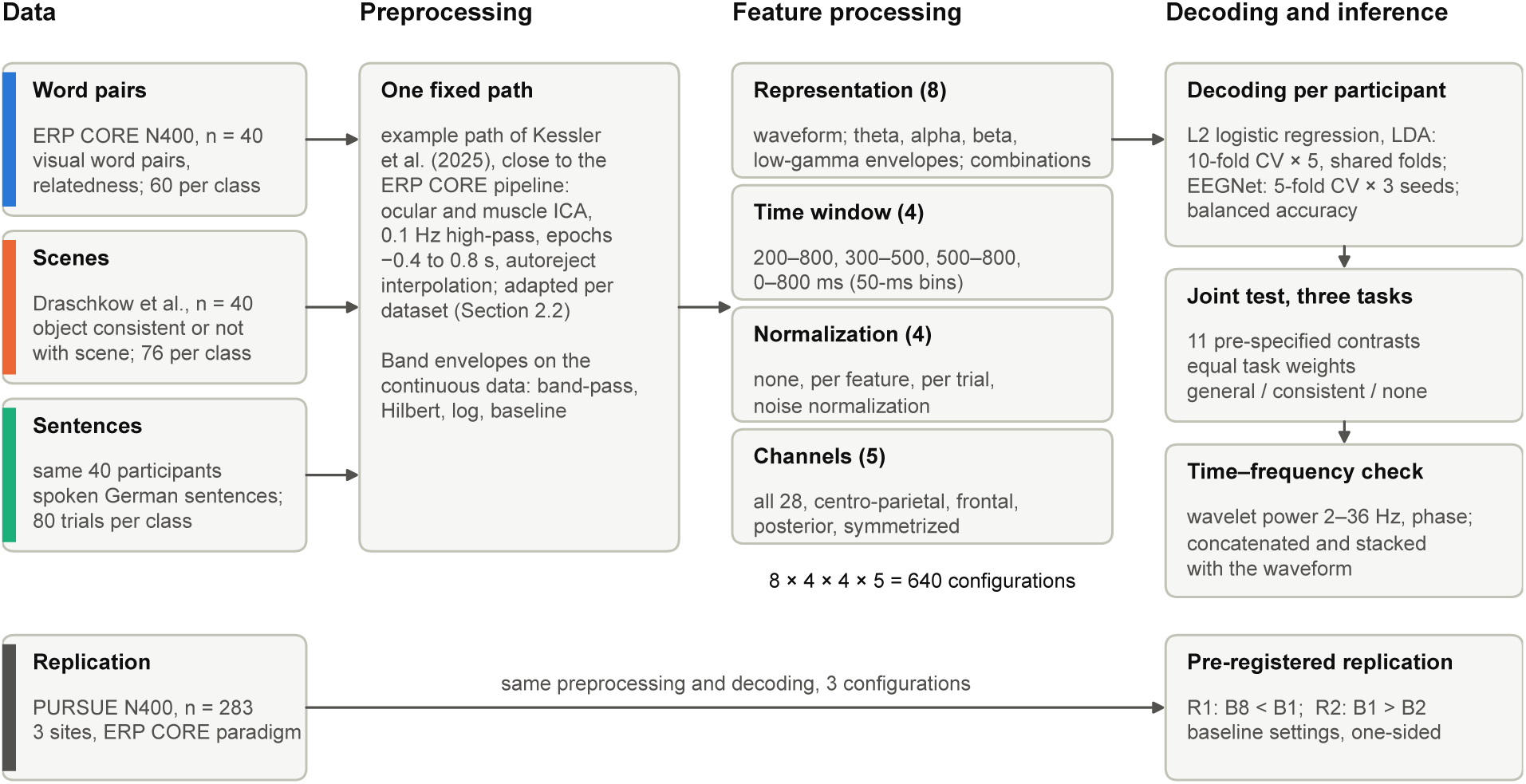
Study design. Three semantic-congruency tasks (visual word pairs, objects in scenes, spoken sentences) were preprocessed with one fixed path, adapted to each dataset where the recordings required it (Section 2.2), and features were computed along four crossed dimensions (representation, time window, normalization, channels; 640 configurations). Each participant was decoded separately with shared cross-validation folds. Eleven pre-specified contrasts were tested jointly across the three tasks; the two most robust conclusions were replicated in a pre-registered study of an independent sample (PURSUE, *n* = 283; three configurations with the same preprocessing and decoding), and a pre-specified time–frequency analysis tested alternative power representations. Numbers in parentheses give the number of levels of each dimension. ICA, independent component analysis; CV, cross-validation.

**Table 2:**
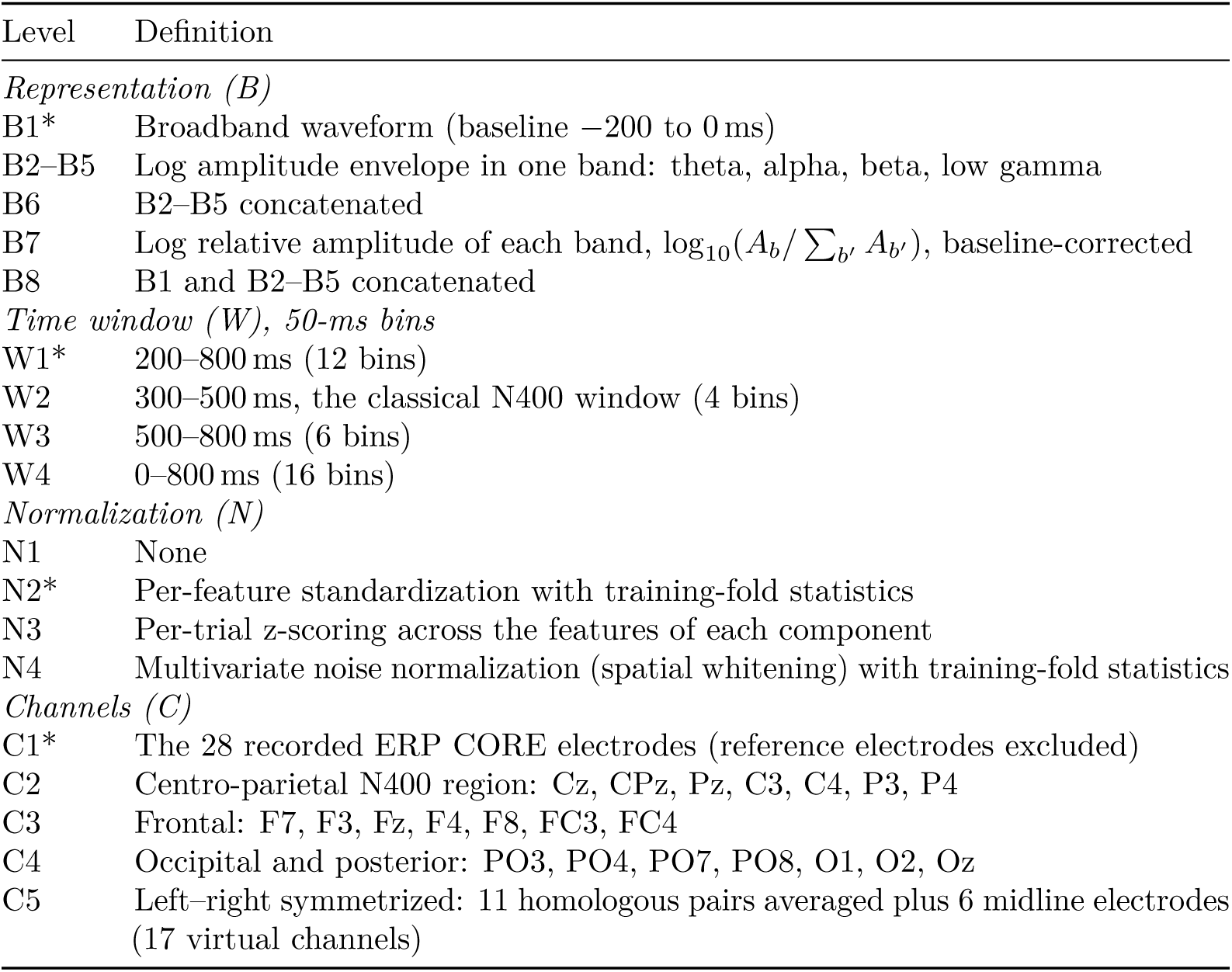
The four feature-processing dimensions. Baseline levels are marked with an asterisk.

| Level | Definition |
|---|---|
| *Representation (B)* |  |
| B1* | Broadband waveform (baseline −200 to 0 ms) |
| B2–B5 | Log amplitude envelope in one band: theta, alpha, beta, low gamma |
| B6 | B2–B5 concatenated |
| B7 | Log relative amplitude of each band, $\log_{10}(A_b/\sum_{b'} A_{b'})$, baseline-corrected |
| B8 | B1 and B2–B5 concatenated |
| *Time window (W), 50-ms bins* |  |
| W1* | 200–800 ms (12 bins) |
| W2 | 300–500 ms, the classical N400 window (4 bins) |
| W3 | 500–800 ms (6 bins) |
| W4 | 0–800 ms (16 bins) |
| *Normalization (N)* |  |
| N1 | None |
| N2* | Per-feature standardization with training-fold statistics |
| N3 | Per-trial z-scoring across the features of each component |
| N4 | Multivariate noise normalization (spatial whitening) with training-fold statistics |
| *Channels (C)* |  |
| C1* | The 28 recorded ERP CORE electrodes (reference electrodes excluded) |
| C2 | Centro-parietal N400 region: Cz, CPz, Pz, C3, C4, P3, P4 |
| C3 | Frontal: F7, F3, Fz, F4, F8, FC3, FC4 |
| C4 | Occipital and posterior: PO3, PO4, PO7, PO8, O1, O2, Oz |
| C5 | Left–right symmetrized: 11 homologous pairs averaged plus 6 midline electrodes (17 virtual channels) |

The representation dimension contrasts the phase-locked broadband signal with band-limited power. Because averaging in 50-ms bins acts as a low-pass filter, B1 effectively carries the signal below about 20 Hz, whereas B2–B5 carry total (phase-locked and non-phase-locked) amplitude in each band. B6 tests whether combining bands helps, B7 removes broadband changes in power and keeps only the spectral composition, and B8 tests whether band power complements the waveform. In multi-component representations each component forms a separate block of features. The time windows test whether the late window (W1 vs. W2) or an early window that should not distinguish the classes (W4 vs. W1) changes performance. Per-feature standardization (N2) equalizes the scale of features across trials; per-trial z-scoring (N3) removes fluctuations in overall amplitude and offset between trials and is computed per component block; multivariate noise normalization (N4; Guggenmos et al., 2018) whitens the channels with the inverse square root of a Ledoit–Wolf shrinkage estimate (Ledoit and Wolf, 2004) of the noise covariance, computed from class-mean-centred training trials for each component. The three regions C2–C4 contain seven electrodes each, so that comparisons between them do not depend on the number of channels; C5 removes all left–right asymmetric information, including lateralized motor activity, while preserving bilaterally distributed effects.

### 2.5 Classification and cross-validation

The main classifier was L2-regularized logistic regression (LR; scikit-learn, lbfgs solver, unpenalized intercept) (Pedregosa et al., 2011). The regularization parameter was chosen in each outer training fold from *{*10*^−^*^3^, 10*^−^*^2^*, . . . ,* 10^2^*}* by stratified 5-fold cross-validation on balanced accuracy. Each participant was modelled separately, with stratified 10-fold cross-validation repeated five times. The 50 outer folds were generated once per participant and shared by all configurations and classifiers, so that all comparisons are paired at the level of folds. All statistics used by the normalizations and by the classifiers were estimated on the outer training fold only. Performance was measured as balanced accuracy averaged over the 50 folds.

Shrinkage linear discriminant analysis (LDA; Ledoit–Wolf shrinkage) served as a second classifier. Because LDA is invariant to per-feature affine transformations, N1 and N2 give identical results; it was run on 480 configurations, with N1 standing in for N2 in all comparisons. EEGNet (Lawhern et al., 2018), implemented in braindecode (Aristimunha et al., 2026; Schirrmeister et al., 2017), was used as a test of generality on the configurations whose input remains a time series (B1–B5 without bin averaging): the baseline and the ten best and three worst of these configurations for logistic regression in each dataset (5-fold cross-validation with three seeds).

### 2.6 Statistical analysis

The unit of analysis was the mean balanced accuracy of a participant in a configuration. Above-chance decoding was tested with one-sided *t*-tests and Wilcoxon signed-rank tests against 0.5, for the baseline and for every configuration (Holm correction across configurations). For individual participants, the baseline was tested with a label-permutation test (1000 permutations).

#### Planned contrasts

Inference was based on eleven contrasts that were specified before any data were analysed (P1–P11; Table 3). Each contrast was computed on participant-level marginal means: for every participant, accuracy was first averaged over the factors not involved in the contrast, and the difference was then taken. For P2 (band combination vs. the best single band), the best band for each participant was chosen from the marginal means of the other participants (leave-one-participant-out), so that the band was never selected on the data of the participant being tested. P11 is an interaction contrast: the contribution of lateralized signal (C1 *−* C5) in the late window (W3) relative to the N400 window (W2).

**Table 3:**
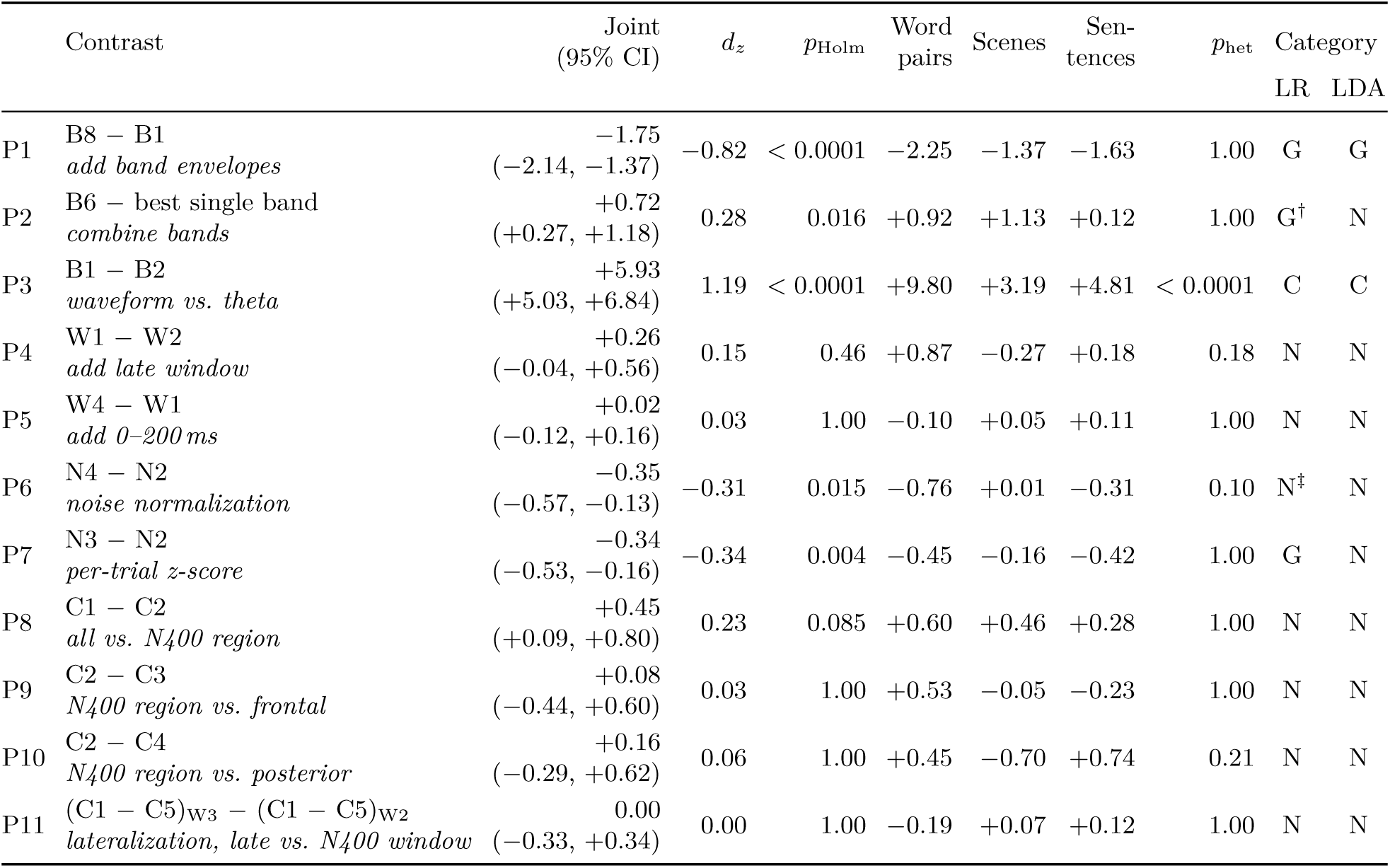
Joint test of the eleven planned contrasts across the three tasks (logistic regression). Estimates in percentage points with 95% confidence intervals; *p*-values Holm-corrected across the eleven contrasts. Category: G, general; C, consistent in direction, varying in size; N, no reliable general effect (no contrast was task-dependent); *p*_het_, heterogeneity test, Holm-corrected. The last two columns give the category for logistic regression (LR) and shrinkage LDA. ^†^Not robust: not significant without one partly flat recording, and for LDA dependent on the preprocessing (Sections 3.3 and 3.7). ^‡^Joint effect significant, but zero in the scene task.

|  | Contrast | Joint (95% CI) | *d*<sub>*z*</sub> | *p*<sub>Holm</sub> | Word pairs | Scenes | Sen-tences | *p*<sub>het</sub> | Category |  |
|---|---|---|---|---|---|---|---|---|---|---|
|  |  |  |  |  |  |  |  |  | LR | LDA |
| P1 | B8 − B1<br>*add band envelopes* | −1.75<br>(−2.14, −1.37) | −0.82 | < 0.0001 | −2.25 | −1.37 | −1.63 | 1.00 | G | G |
| P2 | B6 − best single band<br>*combine bands* | +0.72<br>(+0.27, +1.18) | 0.28 | 0.016 | +0.92 | +1.13 | +0.12 | 1.00 | G<sup>†</sup> | N |
| P3 | B1 − B2<br>*waveform vs. theta* | +5.93<br>(+5.03, +6.84) | 1.19 | < 0.0001 | +9.80 | +3.19 | +4.81 | < 0.0001 | C | C |
| P4 | W1 − W2<br>*add late window* | +0.26<br>(−0.04, +0.56) | 0.15 | 0.46 | +0.87 | −0.27 | +0.18 | 0.18 | N | N |
| P5 | W4 − W1<br>*add 0–200 ms* | +0.02<br>(−0.12, +0.16) | 0.03 | 1.00 | −0.10 | +0.05 | +0.11 | 1.00 | N | N |
| P6 | N4 − N2<br>*noise normalization* | −0.35<br>(−0.57, −0.13) | −0.31 | 0.015 | −0.76 | +0.01 | −0.31 | 0.10 | N<sup>‡</sup> | N |
| P7 | N3 − N2<br>*per-trial z-score* | −0.34<br>(−0.53, −0.16) | −0.34 | 0.004 | −0.45 | −0.16 | −0.42 | 1.00 | G | N |
| P8 | C1 − C2<br>*all vs. N400 region* | +0.45<br>(+0.09, +0.80) | 0.23 | 0.085 | +0.60 | +0.46 | +0.28 | 1.00 | N | N |
| P9 | C2 − C3<br>*N400 region vs. frontal* | +0.08<br>(−0.44, +0.60) | 0.03 | 1.00 | +0.53 | −0.05 | −0.23 | 1.00 | N | N |
| P10 | C2 − C4<br>*N400 region vs. posterior* | +0.16<br>(−0.29, +0.62) | 0.06 | 1.00 | +0.45 | −0.70 | +0.74 | 0.21 | N | N |
| P11 | (C1 − C5)<sub>W3</sub> − (C1 − C5)<sub>W2</sub><br>*lateralization, late vs. N400 window* | 0.00<br>(−0.33, +0.34) | 0.00 | 1.00 | −0.19 | +0.07 | +0.12 | 1.00 | N | N |

#### Joint test across task

The confirmatory test combined the three tasks with equal weight. For each contrast, let *d* denote the participant-level difference. Because the scene and sentence tasks were performed by the same participants, each of these participants contributed the mean of their two differences, *a* = (*d*_scene_ + *d*_sentence_)*/*2. The joint effect,

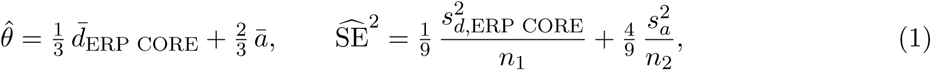

where *n*_1_ and *n*_2_ (40 each) are the numbers of participants in ERP CORE and in the scene and sentence dataset, was tested with a two-sided *t*-test with Welch–Satterthwaite degrees of freedom, and the eleven joint tests were Holm-corrected (Holm, 1979). Heterogeneity across tasks was tested with a two-degree-of-freedom Wald test of the contrasts scene *−* sentence (paired) and ERP CORE *−* mean of the other two (independent), again with Holm correction. Following a pre-specified rule, an effect was classified as *general* if the joint effect was significant, the three task estimates had the same sign and heterogeneity was not significant; as *consistent in direction, varying in size* if the joint effect was significant with the same sign in all tasks but heterogeneity was significant; as *task-dependent* if the signs differed and heterogeneity was significant; and as *no reliable general effect* otherwise. A joint standardized effect *d_z_* was obtained by dividing 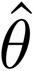 by the root mean square of the three task-wise standard deviations of *d*.

#### Analyses within each dataset

As secondary analyses, each dataset was analysed separately with the planned contrasts (Holm-corrected within the dataset), a four-way repeated-measures analysis of variance with Greenhouse–Geisser correction and generalized *η*^2^ (Olejnik and Algina, 2003; Bakeman, 2005) (Benjamini–Hochberg correction across the four main effects and six two-way interactions; Benjamini and Hochberg, 1995), and an aligned-rank-transform analysis of variance as a non-parametric check (Wobbrock et al., 2011). In a repeated-measures analysis of variance, each effect is tested against its interaction with participants, which corresponds to a maximal by-participant random-effects structure (Barr et al., 2013).

#### Time-resolved decoding

To describe when information becomes available, the baseline settings (B1, N2, logistic regression) were applied to each of the sixteen 50-ms bins between 0 and 800 ms separately, with C1 and C5, on the same folds. Curves were tested against chance with one-sided sign-flip permutation tests with threshold-free cluster enhancement (TFCE; 1024 permutations; Smith and Nichols, 2009), as implemented in MNE-Python (Gramfort et al., 2013).

### 2.7 Pre-registered replication

The two most robust conclusions of the joint test were replicated on the PURSUE dataset in a pre-registered study. The hypotheses, configurations, tests and exclusion criteria were committed to the study repository before the EEG data were downloaded; the pre-registration document is available in the public repository (see Data and code availability). Three configurations were decoded with the same procedure: the baseline (B1-W1-N2-C1), the waveform plus band envelopes (B8-W1-N2-C1) and the theta envelope (B2-W1-N2-C1). R1 predicted lower accuracy for B8 than for B1 (corresponding to P1), and R2 higher accuracy for B1 than for B2 (corresponding to P3). Both were tested with one-sided paired *t*-tests, Holm-corrected, with Wilcoxon tests as a check; replication was declared if both were significant in the predicted direction. Recordings without EEG data were excluded in advance (three participants), and participants with fewer than 30 trials in either class after preprocessing were excluded (one participant). One recording used a different 32-channel montage that lacked ten of the pre-registered electrodes and was excluded as a deviation from the pre-registration, leaving 283 participants. LDA, results per site and results without the post-preprocessing exclusion were pre-registered as secondary analyses.

### 2.8 Time–frequency sensitivity analysis

A pre-specified sensitivity analysis tested whether the conclusions about band power reflected a representation that was too coarse (four broad bands, no delta, amplitude only) or a concatenation that disadvantages power by forcing many power features to share one regularization parameter with fewer waveform features. The continuous data were decomposed with complex Morlet wavelets at 12 log-spaced frequencies between 2 and 36 Hz (3 to 10 cycles), epoched and interpolated as in Section 2.3. With the other dimensions at baseline and logistic regression only, five representations were decoded on the main folds: log wavelet amplitude at all 12 frequencies (T1) and at the six frequencies up to 7.4 Hz (T2), the cosine and sine of the phase at these six frequencies (T3), the waveform concatenated with T1 (T4), and the stacked combination of separate waveform and T1 models (T5). For stacking (Wolpert, 1992), out-of-sample decision values of the two base models were obtained by 5-fold cross-fitting within each outer training fold with the regularization parameter fixed to the value selected for that fold, standardized, and combined by a logistic-regression meta-model. Six comparisons (F1: T4 *−* B1; F2: T5 *−* B1; F3: T1 *−* B6; F4: T1 *−* B1; F5: T2 *−* T1; F6: T3 *−* B1) were tested with the joint test (Holm across the six), F1 and F2 being the primary comparisons. The interpretation of the possible outcomes was fixed before the analysis was run (details in Supplementary Section S9).

### 2.9 Robustness analyses

Three further analyses checked that the conclusions did not depend on analysis choices. First, all logistic-regression configurations of the three tasks were re-run on the same folds with an extended regularization grid (10*^−^*^5^ to 10^4^). Second, the full analysis of the scene and sentence tasks was re-run with the preprocessing specified before the pilot check (Section 2.2). Third, the joint test was repeated without the one sentence-task participant whose recording was partly flat (Supplementary Section S7).

### 2.10 Degree of pre-specification

The eleven planned contrasts and their analysis were fixed before any data were analysed. The joint test and its classification rule were specified after the ERP CORE results were known but before any decoding of the scene and sentence tasks. The time–frequency analysis was specified before it was run, and the replication was pre-registered. All other analyses are labelled as secondary or exploratory.

### 2.11 Software

Analyses used Python 3.11 with MNE-Python 1.13.2 (Gramfort et al., 2013; Larson et al., 2026), autoreject 0.5 (Jas et al., 2017), python-picard 0.8 (Ablin et al., 2018a,b), scikit-learn 1.8 (Pedregosa et al., 2011), braindecode 1.8.1 (Aristimunha et al., 2026; Schirrmeister et al., 2017), NumPy, SciPy and pandas. The full list of package versions and fixed random seeds is part of the code repository (see Data and code availability).

## 3 Results

Differences in balanced accuracy are reported in percentage points (pp; 1 pp = 0.01). Unless stated otherwise, results refer to logistic regression.

### 3.1 Data retention and single-trial decodability

Preprocessing retained 98.8% of the trials in ERP CORE (4743 of 4800), 99.9% in the scene task (6075 of 6080), 89.1% in the sentence task (5704 of 6401; 84–160 trials per participant) and 99.4% in PURSUE (Supplementary Section S1). With the baseline configuration, semantic congruency was decodable from single trials in all three tasks (all *p <* 10*^−^*^8^; Table 4, Fig. 3A), but accuracy depended strongly on the task: 66.3% for visual word pairs, 57.0% for spoken sentences and 55.3% for objects in scenes. Correspondingly, permutation tests were significant for 33, 24 and 16 of the 40 participants (two expected by chance). The order of the tasks followed the size of the N400 effect over the centro-parietal region (300–500 ms: *−*5.3, *−*3.0 and *−*1.0 µV), and in ERP CORE the individual N400 effect explained about half of the between-participant variance in accuracy (*r* = *−*0.73; sentences *r* = *−*0.35; scenes *r* = *−*0.12, n.s.; exploratory, Supplementary Section S8).

**Figure 3:**
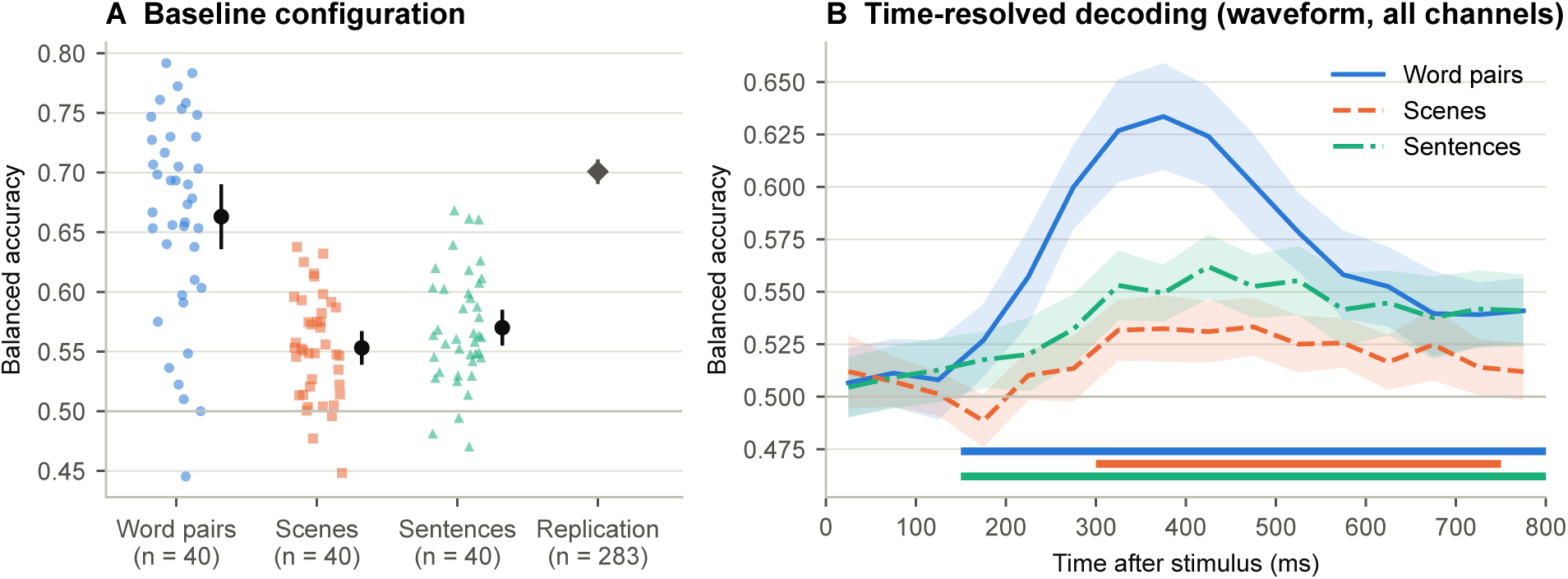
Single-trial decodability. (A) Balanced accuracy of the baseline configuration for each participant in the three tasks (dots) with the mean and 95% confidence interval (black), and the mean and 95% confidence interval in the replication sample (grey diamond). (B) Time-resolved decoding in 50-ms bins (waveform, all channels, logistic regression); shaded areas, 95% confidence intervals; horizontal bars below the curves (from top to bottom: word pairs, scenes, sentences), bins above chance in one-sided permutation tests with threshold-free cluster enhancement.

**Table 4:**
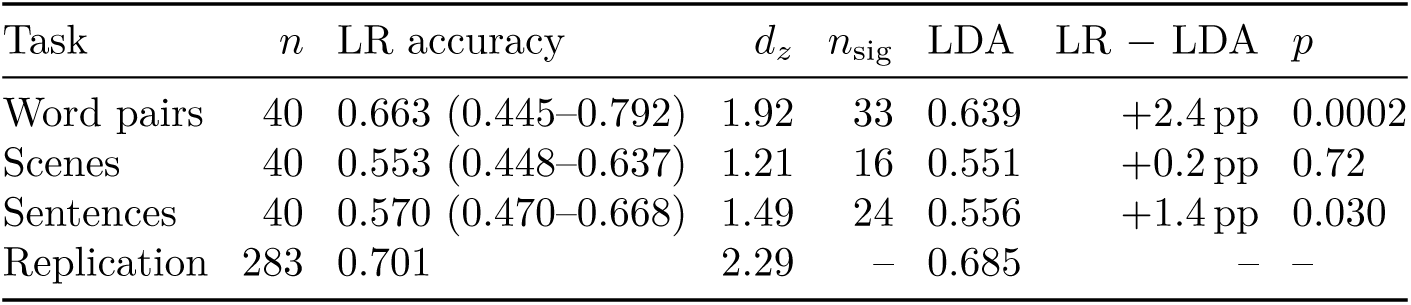
Baseline configuration (B1-W1-N2-C1). Mean balanced accuracy (range across participants), standardized effect against chance, number of participants with a significant permutation test (*n*_sig_), and the difference between logistic regression (LR) and shrinkage LDA, with the *p*-value of a paired *t*-test; –, not computed.

| Task | *n* | LR accuracy | *d<sub>z</sub>* | *n*<sub>sig</sub> | LDA | LR − LDA | *p* |
|---|---|---|---|---|---|---|---|
| Word pairs | 40 | 0.663 (0.445–0.792) | 1.92 | 33 | 0.639 | +2.4 pp | 0.0002 |
| Scenes | 40 | 0.553 (0.448–0.637) | 1.21 | 16 | 0.551 | +0.2 pp | 0.72 |
| Sentences | 40 | 0.570 (0.470–0.668) | 1.49 | 24 | 0.556 | +1.4 pp | 0.030 |
| Replication | 283 | 0.701 | 2.29 | – | 0.685 | – | – |

Time-resolved decoding showed the same task ordering (Fig. 3B). Decoding became significant at 150–200 ms for word pairs and sentences and at 300–350 ms for scenes, and peaked within the N400 window in all tasks (word pairs 350–400 ms, 63.4%; sentences 400–450 ms, 56.2%; scenes 450–500 ms, 53.3%), declining slowly thereafter and remaining above chance until 800 ms for word pairs and sentences and until 750 ms for scenes.

### 3.2 Representation is the dominant dimension

In every task, the signal representation had by far the largest effect on accuracy (Table 5, Fig. 4). Its generalized *η*^2^ averaged 0.165 across tasks, an order of magnitude above the next effects (the representation *×* window interaction, 0.016; the time window, 0.010), and the main effect of representation was significant in all tasks (*p <* 0.0001; confirmed by the aligned-rank-transform analysis; Supplementary Section S3). Normalization and channel set had effects with generalized *η*^2^ *≤* 0.008. The waveform-based representations (B1, B8) were the only ones that performed well: marginal means were 61.5%, 53.3% and 54.8% for B1 and 59.3%, 52.0% and 53.2% for B8 (word pairs, scenes, sentences), whereas the representations based on envelopes alone reached 50.9–52.7% in ERP CORE and 49.6–50.4%, i.e. chance, in the other two tasks. Accordingly, of the configurations decoded above chance after Holm correction (173, 65 and 109 of 640), all of those in the scene and sentence tasks used B1 or B8.

**Figure 4:**
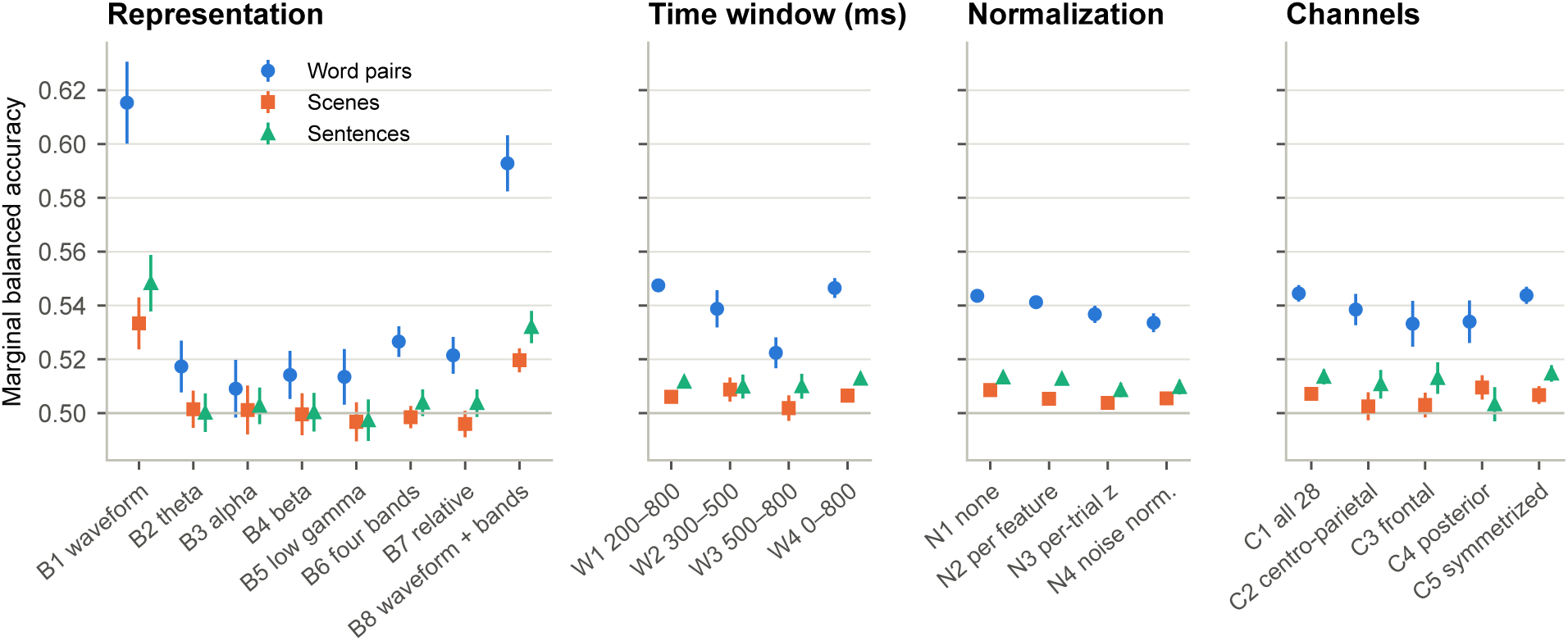
Marginal mean balanced accuracy for each level of the four dimensions in the three tasks (logistic regression; each participant’s accuracy averaged over the other three dimensions). Error bars, within-participant 95% confidence intervals (Cousineau–Morey). All panels share the vertical axis; the horizontal line marks chance. Levels as in Table 2.

**Table 5:** Generalized *η*^2^ of the largest effects in the four-way repeated-measures analysis of variance (logistic regression).

| Effect | Word pairs | Scenes | Sentences | Mean |
|---|---|---|---|---|
| Representation (B) | 0.289 | 0.079 | 0.127 | 0.165 |
| B × W | 0.035 | 0.011 | 0.002 | 0.016 |
| Time window (W) | 0.027 | 0.003 | 0.001 | 0.010 |
| B × C | 0.005 | 0.008 | 0.007 | 0.007 |
| Channels (C) | 0.006 | 0.004 | 0.008 | 0.006 |
| B × N | 0.007 | 0.004 | 0.004 | 0.005 |
| Normalization (N) | 0.004 | 0.002 | 0.002 | 0.003 |

### 3.3 Joint test across tasks

The joint test identified two robust conclusions about the representation (Table 3, Fig. 5). The waveform outperformed the theta envelope by 5.9 pp (P3; *d_z_* = 1.19); the effect had the same sign in all tasks but differed in size (largest for word pairs, 9.8 pp), and was therefore classified as consistent in direction but varying in size. Adding the four band envelopes to the waveform lowered accuracy by 1.8 pp (P1; *d_z_* = *−*0.82), significantly so in each task and without heterogeneity, and was classified as general. Both conclusions held for LDA (P1 *−*2.6 pp; P3 +6.0 pp; Supplementary Section S4), with the extended regularization grid, with the originally specified preprocessing and without the partly flat recording described below (Section 3.7).

**Figure 5:**
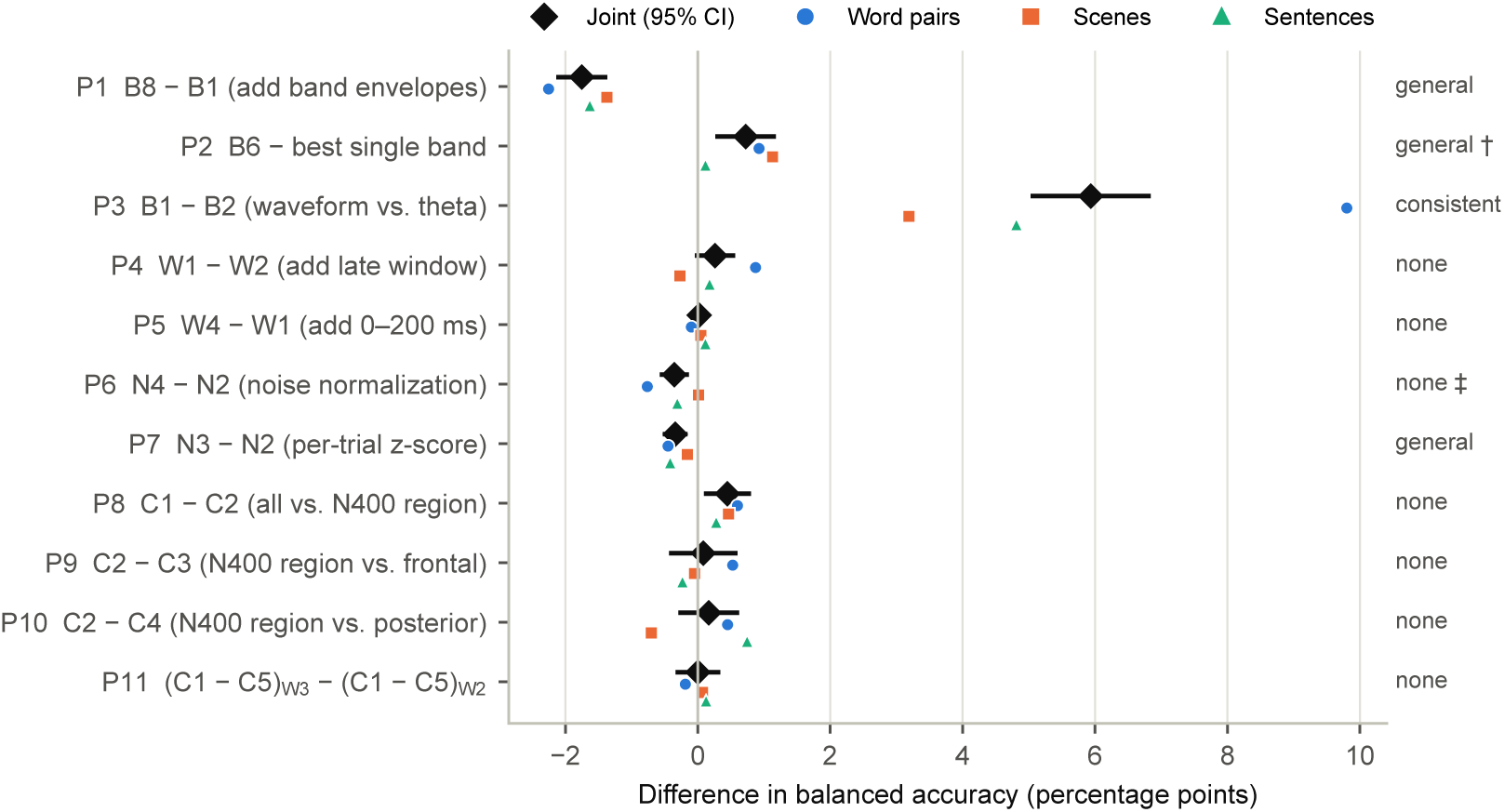
Joint test of the planned contrasts (logistic regression). For each contrast, the joint estimate with its 95% confidence interval (black diamond) and the estimates of the three tasks (coloured symbols); the pre-specified category is given on the right (general; consistent = consistent in direction but varying in size; none = no reliable general effect). **^†^**Not robust to one partly flat recording or, for LDA, to the preprocessing. ^‡^Joint effect significant, but zero in the scene task.

Per-trial z-scoring was slightly worse than per-feature standardization in all tasks (P7, *−*0.34 pp, general), but only for logistic regression. Multivariate noise normalization was also slightly worse than per-feature standardization overall (P6, *−*0.35 pp, *p*_Holm_ = 0.015), but not in the scene task (+0.01 pp), so that, by the pre-specified rule, it did not count as a general effect. Combining the four bands was slightly better than the best single band (P2, +0.72 pp), but this effect depended on a single participant whose sentence recording was partly flat: without this participant it was no longer significant (*p*_Holm_ = 0.38; Supplementary Section S7), and for LDA it was significant only with the originally specified preprocessing (Section 3.7). None of the contrasts on time windows (P4, P5, P11) or channel sets (P8–P10) showed a reliable general effect; in particular, the full set of 28 channels did not differ reliably from the seven centro-parietal electrodes (P8, +0.45 pp). No contrast was task-dependent.

### 3.4 Pre-registered replication

Both pre-registered predictions were confirmed in the independent PURSUE sample (*n* = 283; Fig. 6). Adding the band envelopes to the waveform lowered accuracy (R1: *−*3.6 pp, 95% CI *−*4.1 to *−*3.0, *d_z_*= *−*0.76, *p*_Holm_ *<* 10*^−^*^15^), and the waveform outperformed the theta envelope (R2: +15.0 pp, 95% CI +13.9 to +16.0, *d_z_*= 1.64, *p*_Holm_ *<* 10*^−^*^15^); the baseline reached 70.1% (*d_z_*= 2.29). The replication thus succeeded. Both effects were comparable to or larger than the corresponding effects with the other dimensions at baseline in ERP CORE (*−*3.7 and +13.5 pp), held for LDA (*d_z_* = *−*0.61 and 1.50), were virtually unchanged without the post-preprocessing exclusion (*n* = 284), and held at each of the three sites (R1 *d_z_* from *−*0.70 to *−*0.86; R2 from 1.56 to 1.74).

**Figure 6:**
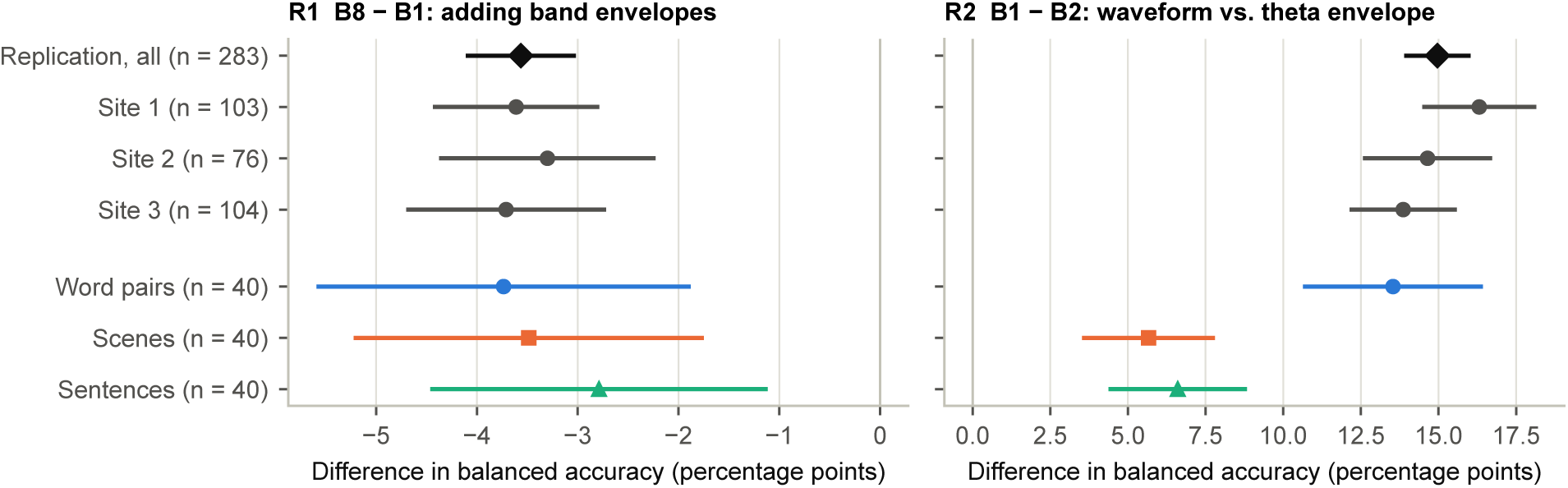
Pre-registered replication in PURSUE. Mean differences in balanced accuracy for R1 (B8 *−* B1) and R2 (B1 *−* B2) in the whole sample (*n* = 283, black diamond) and at each site (grey), with the same differences in the three original tasks for comparison (other dimensions at baseline; logistic regression). Horizontal lines, 95% confidence intervals. Sites 1–3 are the groups of participant codes 1xxx, 2xxx and 3xxx in the dataset, which does not state which institution each group belongs to.

### 3.5 Richer power representations do not add information

The time–frequency analysis confirmed and strengthened the conclusions about band power (Table 6, Fig. 7). The wavelet representation was indeed better than the original four bands (F3, +1.5 pp, general), but used alone it remained far below the waveform (F4, *−*6.2 pp). Concatenating it with the waveform lowered accuracy in all tasks (F1, *−*3.4 pp, general), and so did stacking separate waveform and power models (F2, *−*1.4 pp, general), even though the meta-model assigned power a positive weight in most folds (median share 0.13–0.22): in data of this size, power provided no information that was usable beyond the waveform. Frequencies above 8 Hz added nothing (F5). The phase of the low frequencies came close to the waveform in ERP CORE (65.1% vs. 66.3%) and fell further behind in the other two tasks (F6), consistent with the information residing in phase-locked low-frequency activity.

**Figure 7:**
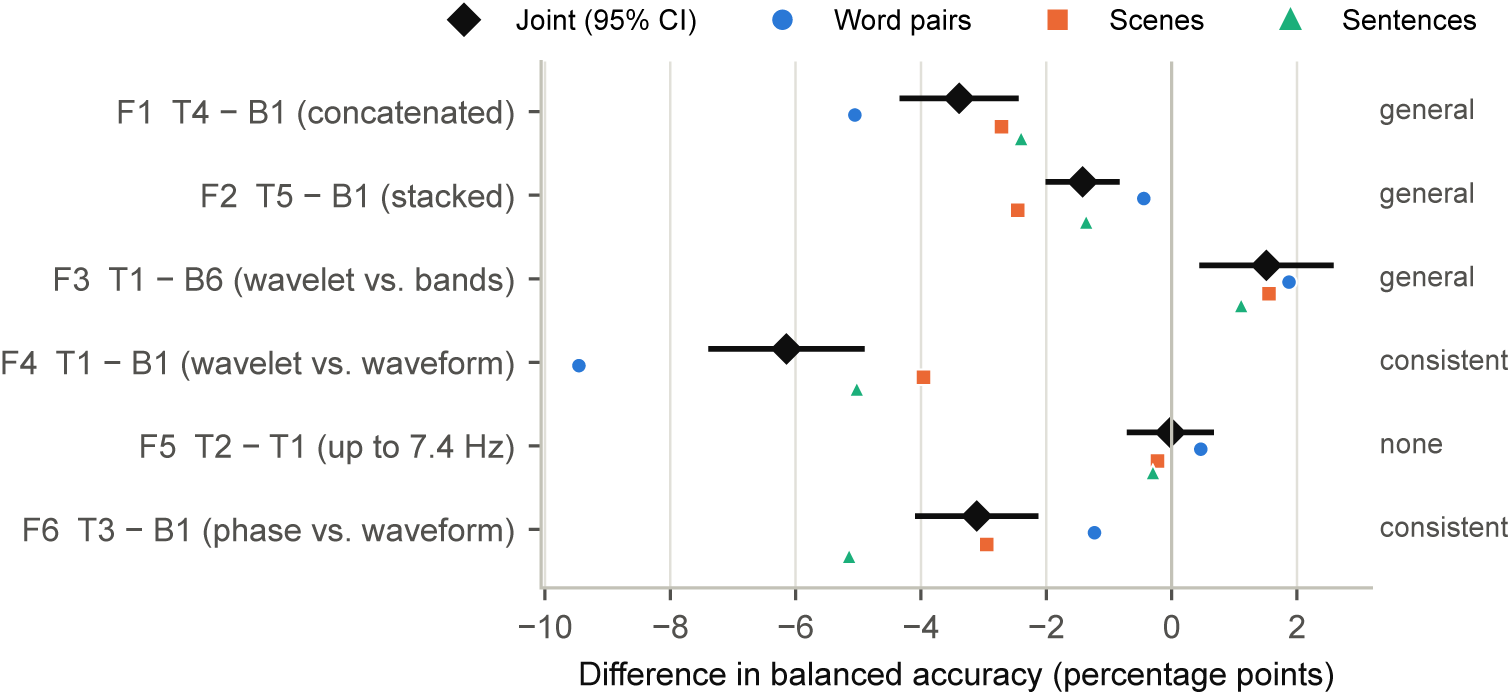
Time–frequency sensitivity analysis (logistic regression). Joint estimate with its 95% confidence interval (black diamond) and the estimates of the three tasks (coloured symbols) for the six pre-specified comparisons; categories as in Fig. 5. T1, log wavelet amplitude at 12 frequencies (2–36 Hz); T2, the same up to 7.4 Hz; T3, phase up to 7.4 Hz; T4, waveform and T1 concatenated; T5, waveform and T1 models stacked.

**Table 6:**
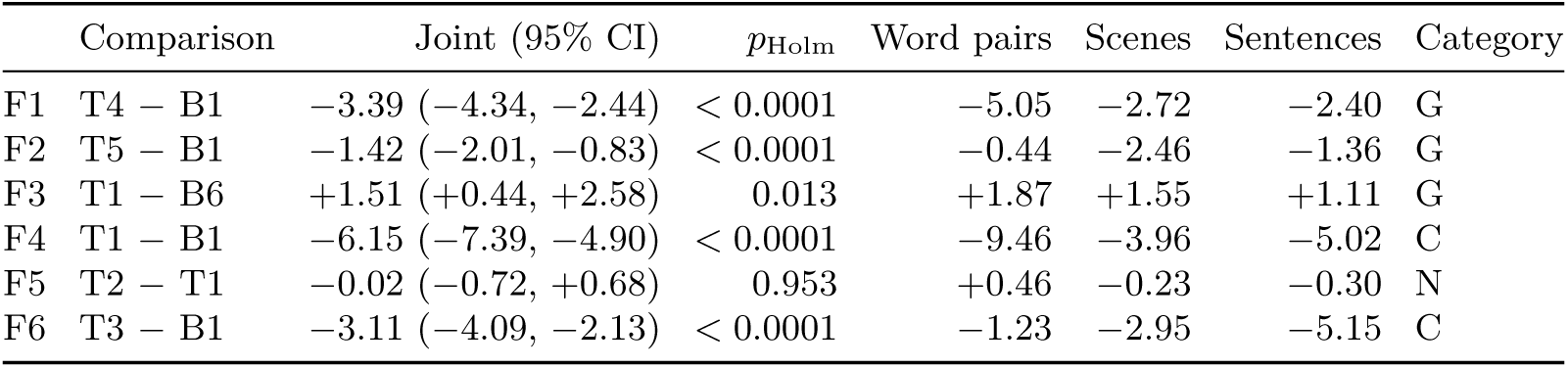
Time–frequency sensitivity analysis: joint test across the three tasks (logistic regression; percentage points; *p*-values Holm-corrected across the six comparisons). T1, log wavelet amplitude at 12 frequencies (2–36 Hz); T2, the same up to 7.4 Hz; T3, phase up to 7.4 Hz; T4, waveform and T1 concatenated; T5, waveform and T1 models stacked. Categories as in Table 3.

### 3.6 The conventional configuration lies on a plateau

Averaged over the three tasks, the baseline configuration was the best of all 640 (59.5%). It ranked first for word pairs and sentences and second for scenes, 0.1 pp behind the same configuration with per-trial z-scoring. Nine of the ten best configurations used the waveform alone and one the waveform with envelopes (58.5–59.5%), forming a plateau whose internal order was not stable across tasks or classifiers. Configuration means were strongly correlated across tasks (Pearson *r* = 0.83, 0.89 and 0.76 for word pairs and scenes, word pairs and sentences, and scenes and sentences); rank correlations were lower (0.40–0.68) because the order of the many near-chance envelope configurations is dominated by noise. In the 40 participants who performed both the scene and the sentence task, baseline accuracies were not significantly correlated (*r* = 0.25, *p* = 0.12).

### 3.7 Classifiers and robustness

Logistic regression outperformed LDA at baseline for word pairs and sentences and matched it for scenes (Table 4). In all tasks, LDA was slightly better with few features and worse with many (correlation of the difference with the logarithm of the number of features among waveform configurations, *r* = *−*0.42, *−*0.48 and *−*0.37), as expected from its need to estimate the feature covariance. EEGNet was on average 1.0–1.2 pp below logistic regression on the waveform configurations and exceeded it in none of them (Supplementary Section S5).

The extended regularization grid did not improve accuracy (baseline *−*0.2 to *−*0.5 pp; correlation of the 640 configuration means *≥* 0.987), left the significance of all 33 within-dataset planned contrasts unchanged and changed no joint category except that of P6, which changed from no reliable general effect to general because its near-zero estimate in the scene task changed sign (+0.01 to *−*0.04 pp; Supplementary Section S6). Re-running the scene and sentence tasks with the originally specified preprocessing (all 60 EEG channels, muscle independent component analysis and interpolation in both tasks) did not change baseline accuracy significantly (scenes *−*0.2 pp, *p* = 0.51; sentences *−*0.9 pp, *p* = 0.12; correlation of the configuration means 0.99 and 0.91) and left all eleven joint categories unchanged for logistic regression (P1 *−*1.7 pp, P3 +5.8 pp, P7 *−*0.3 pp; Supplementary Section S2). For LDA, one category changed: the benefit of combining bands (P2) became significant (+0.9 pp, *p*_Holm_ = 0.003; main analysis +0.7 pp, *p*_Holm_ = 0.11), so that its classification depends on the preprocessing as well as on a single participant.

## 4 Discussion

We compared 640 feature-processing configurations for single-trial decoding of semantic congruency in three tasks that differ in modality and in the kind of semantic context. One dimension stood out. The signal representation had by far the largest effect in every task, the broadband waveform outperformed band-limited amplitude envelopes, and adding band power to the waveform reduced accuracy in all three tasks. Both conclusions replicated in a pre-registered study of an independent sample of 283 participants, and neither a finer wavelet decomposition including delta nor a stacking scheme that does not force power and waveform features to share one regularization parameter allowed power to add information. By contrast, the time window, the normalization and the channel set had small or no general effects, and the conventional configuration—waveform, 200–800 ms, per-feature standardization, all channels—lay on a plateau of near-optimal configurations.

That the waveform outperforms a single frequency band is in line with common practice in ERP research and was expected. Less expected were that adding band power lowered accuracy rather than leaving it unchanged, that this held across modalities and kinds of semantic context and in a pre-registered replication, and that the other feature-processing choices mattered so little. These findings are relevant beyond the ERP community, because EEG-to-text research uses band power as its default representation.

### 4.1 Why the waveform carries the information

Several observations indicate that the decodable semantic information is carried by phase-locked activity at low frequencies. The waveform, which after averaging in 50-ms bins contains mainly the signal below about 20 Hz, was the best representation in all tasks. The phase of the wavelet coefficients below 8 Hz came close to the waveform in ERP CORE, whereas power at the same frequencies did not, and power above 8 Hz added nothing. In an exploratory analysis of ERP CORE, the condition differences in the band envelopes over the centro-parietal region were all within 4% of the amplitude, whereas the N400 effect in the waveform amounted to *−*5.3 µV (a single-trial effect size of *d* = *−*0.62), and the size of each participant’s N400 effect explained about half of the between-participant variance in accuracy. Band envelopes measure the total amplitude in a band and discard its phase; the part of an event-related power change that stems from the evoked potential is captured more directly by the waveform, and the remainder appears to be small relative to the variability of power across trials.

The advantage of the waveform did not depend on the window over which the features were computed. In each of the four time windows, from the 200-ms N400 window to the full 800 ms after stimulus onset, every single-band envelope remained within 2.3 percentage points of chance in word pairs and within 1.3 points in the other two tasks, whereas the waveform reached 56.0–63.9% and 52.0–55.2%, respectively; adding the envelopes to the waveform lowered accuracy in every window and task (Supplementary Section S3). Nor did the result depend on the length of the window over which power was estimated: the wavelets of the time–frequency analysis spanned 3 to 10 cycles, from 1.5 s at 2 Hz to 0.3 s at 36 Hz, and wavelet power, although better than the four band envelopes, also remained far below the waveform (F3, F4).

Adding power to the waveform was not merely useless but harmful. With concatenation, the 1344 envelope features of the baseline window outnumber the 336 waveform features four to one and share one regularization parameter with them, so that noise in the power features dilutes the waveform. That this is not the whole explanation is shown by the stacking analysis: when separate models were fitted to the waveform and to wavelet power and combined by a meta-model, power still received a positive weight in most folds but reduced accuracy on the test trials. With 60–80 trials per class, the power features therefore contributed variance that could be fitted but not generalized.

These results do not contradict the literature on oscillatory correlates of semantic processing. Semantic violations are accompanied by changes in theta and gamma power in trial averages (Hald et al., 2006); in our exploratory analysis, late theta amplitude also increased, but the change was small, as was a late decrease in beta amplitude. Differences between condition averages need not translate into single-trial discriminability, and part of an event-related power change can reflect the evoked component itself. Our conclusion is also specific to semantic congruency. In other domains, power and potentials carry different information: for example, alpha power tracks the attended location but carries little or no information about the orientation held in working memory (Bae and Luck, 2018). We do not claim that band power is uninformative in general, only that it does not add to the waveform for the semantic effects studied here.

### 4.2 Implications for semantic decoding and EEG-to-text research

EEG-to-text studies typically use band-power features aggregated over the fixations on each word (Hollenstein et al., 2018; Wang and Ji, 2022). Our results suggest that such a representation discards much of the single-trial semantic information available in the EEG. Together with the report that some of these models perform similarly when the EEG input is replaced by noise (Jo et al., 2024), this points to the input representation as one candidate explanation. We did not test naturalistic reading, fixation-locked signals or sentence-level decoding, so this is an implication rather than a direct test; decoding the same corpora with waveform features would be a straightforward next step. For N400-based brain–computer interfaces (Dijkstra et al., 2020), the results support waveform features with a regularized linear classifier. EEGNet did not outperform logistic regression in any task, and in single-trial N400 decoding for augmented reality, shrinkage LDA, support vector machines and EEGNet performed comparably (Wimmer et al., 2024): with data of this size, the limiting factor appears to be the information in the features rather than the flexibility of the classifier.

### 4.3 Implications for decoding methodology

Recent methodological work has evaluated preprocessing and analysis parameters, mostly one at a time (Zhang et al., 2024; Zhang and Luck, 2025; Zhang et al., 2026a,b; Qin et al., 2026) and, for preprocessing, in a multiverse of all combinations (Kessler et al., 2025). These studies used different classifiers and measures, often with averages of several trials, so their effect sizes cannot be compared directly with ours. Within the choices we examined, however, the representation was the only one with a large effect, and the remaining choices mattered little, both individually and in interaction. Two practical recommendations follow. First, the representation should be chosen deliberately, and band envelopes should not replace or be added to the waveform for semantic decoding. Second, once a sensible configuration has been chosen, little is gained by optimizing the time window, the normalization or the channel set: the conventional configuration was the best on average across tasks, the order of configurations within the plateau was not stable across tasks or classifiers, and searching within the plateau mainly adds analytic flexibility.

Some expectations from other fields were not supported. Multivariate noise normalization, recommended for MEG decoding (Guggenmos et al., 2018), was at best equal to per-feature standardization (scene task) and slightly worse overall, in line with a report from fMRI that its benefit is unreliable (Ritchie et al., 2021). Per-trial z-scoring was slightly but consistently harmful, consistent with the finding that sample-wise spatial normalization helps between-participant but not within-participant decoding (Qin et al., 2026). The full set of 28 channels was not reliably better than the seven centro-parietal electrodes, and the three regions did not differ reliably, which agrees with a report that the benefit of more electrodes depends on the task (Hou et al., 2026) and with the finding that spatial dimensionality reduction does not improve decoding (Zhang et al., 2024).

### 4.4 Differences between tasks

Accuracy differed considerably between tasks and followed the size of the N400 effect: 66% for visual word pairs with the largest effect, 55% for scenes with a group-average effect of only 1 µV. In the scene task the individual N400 effect did not predict accuracy and decoding peaked later (450–500 ms), suggesting that the classifier exploited activity beyond the centro-parietal N400. The tasks differ in modality, task demands and participants, so the sources of these differences cannot be separated here. The representation *×* time window interaction was notable only in ERP CORE, which was also the only task with a speeded judgement. What matters for our conclusions is that the effects of representation had the same direction in all three tasks despite these differences, which is what the joint test was designed to establish.

### 4.5 Limitations

Five limitations should be noted. First, the benefit of combining frequency bands (P2) depended on a single participant and, for LDA, on the preprocessing, and should not be taken as a conclusion. Second, EEGNet was evaluated on 13 time-series configurations per task, not on the full design. Third, with 60–80 trials per class, small amounts of information in power that is complementary to the waveform may not be usable; whether power adds information with substantially more trials, for example in multi-session recordings, remains open. Fourth, the time–frequency analysis was run with logistic regression only. Fifth, the features were always averaged in 50-ms bins; other bin widths were not compared.

## 5 Conclusions

For single-trial decoding of semantic congruency from EEG, the choice of signal representation matters far more than the time window, the normalization or the channel set. The broadband waveform carries the information; band-limited power does not add to it and, when added, reduces accuracy. These conclusions hold across visual word pairs, scenes and spoken sentences, were replicated in a pre-registered study of 283 participants, and survive a finer time–frequency decomposition and a fairer way of combining power with the waveform. For semantic decoding, we recommend waveform features with a broad post-stimulus window and per-feature standardization, and caution against band-power representations.

## Data and code availability

All EEG data are openly available: ERP CORE at https://doi.org/10.18115/D5JW4R (Luck, 2020), the scene and sentence data at https://doi.org/10.5281/zenodo.1421347 (CC BY 4.0; Draschkow et al., 2018a), and PURSUE at https://doi.org/10.82901/nemar.on007052 (CC0; Couperus et al., 2026). The analysis code, the pre-registration document of the PURSUE replication and the participant-level decoding results are available at https://github.com/LaurentiusY/ERP-N400-Feature-Processing-Study-Public.

## CRediT authorship contribution statement

**Laurence Yang:** Conceptualization, Methodology, Software, Formal analysis, Visualization, Writing – original draft, Writing – review & editing. **Milad Lankarany:** Supervision, Writing – review & editing.

## Declaration of generative AI and AI-assisted technologies in the manuscript preparation process

During the preparation of this work the first author used a large language model (Claude, Anthropic) to help write the analysis code and prepare the manuscript. The first author checked all code, results and text, and both authors take full responsibility for the content of this manuscript.

## Acknowledgements

We thank the authors of ERP CORE, of the scene and sentence dataset and of the PURSUE dataset for making their data openly available. Computing resources were provided by the Digital Research Alliance of Canada.

## References

Ablin, P., Cardoso, J.F., Gramfort, A., 2018a. Faster ICA under orthogonal constraint, in: 2018 IEEE International Conference on Acoustics, Speech and Signal Processing (ICASSP), pp. 4464–4468. doi:10.1109/ICASSP.2018.8461662.

Ablin, P., Cardoso, J.F., Gramfort, A., 2018b. Faster independent component analysis by preconditioning with Hessian approximations. IEEE Transactions on Signal Processing 66, 4040–4049. doi:10.1109/TSP.2018.2844203.

Aristimunha, B., Guetschel, P., Wimpff, M., Gemein, L., Rommel, C., Banville, H., Sliwowski, M., Wilson, D., Brandt, S., Gnassounou, T., Paillard, J., Srivastava, A., Junqueira Lopes, B., Sedlar, S., Tayal, S., Truong, Y., Moreau, T., Chevallier, S., Gramfort, A., Schirrmeister, R.T., 2026. Braindecode: Toolbox for decoding raw electrophysiological brain data with deep learning models. Zenodo, version 1.8.1. doi:10.5281/zenodo.22209839.

Bae, G.Y., Luck, S.J., 2018. Dissociable decoding of spatial attention and working memory from EEG oscillations and sustained potentials. Journal of Neuroscience 38, 409–422. doi:10.1523/JNEUROSCI.2860-17.2017.

Bakeman, R., 2005. Recommended effect size statistics for repeated measures designs. Behavior Research Methods 37, 379–384. doi:10.3758/BF03192707.

Barr, D.J., Levy, R., Scheepers, C., Tily, H.J., 2013. Random effects structure for confirmatory hypothesis testing: Keep it maximal. Journal of Memory and Language 68, 255–278. doi:10.1016/j.jml.2012.11.001.

Benjamini, Y., Hochberg, Y., 1995. Controlling the false discovery rate: A practical and powerful approach to multiple testing. Journal of the Royal Statistical Society: Series B (Methodological) 57, 289–300. doi:10.1111/j.2517-6161.1995.tb02031.x.

Blankertz, B., Lemm, S., Treder, M., Haufe, S., Müller, K.R., 2011. Single-trial analysis and classification of ERP components—A tutorial. NeuroImage 56, 814–825. doi:10.1016/j.neuroimage.2010.06.048.

Carrasco, C.D., Bahle, B., Simmons, A.M., Luck, S.J., 2024. Using multivariate pattern analysis to increase effect sizes for event-related potential analyses. Psychophysiology 61, e14570. doi:10.1111/psyp.14570.

Chen, L., Cichy, R.M., Kaiser, D., 2022. Semantic scene-object consistency modulates N300/400 EEG components, but does not automatically facilitate object representations. Cerebral Cortex 32, 3553–3567. doi:10.1093/cercor/bhab433.

Couperus, J.W., Bukach, C.M., Reed, C.L., 2026. PURSUE N400 word processing. NEMAR (Neuroelectromagnetic Data Archive and Tools Resource), dataset on007052, version 1.0.0. doi:10.82901/nemar.on007052.

Couperus, J.W., Lydic, K.O., Hollis, J.E., Roy, J.L., Lowe, A.R., Bukach, C.M., Reed, C.L., 2021. Individual differences in working memory and the N2pc. Frontiers in Human Neuroscience 15, 620413. doi:10.3389/fnhum.2021.620413.

Dijkstra, K.V., Farquhar, J.D.R., Desain, P.W.M., 2020. The N400 for brain computer interfacing: Complexities and opportunities. Journal of Neural Engineering 17, 022001. doi:10.1088/1741-2552/ab702e.

Draschkow, D., Heikel, E., Võ, M.L.H., Fiebach, C.J., Sassenhagen, J., 2018a. Electroencephalography dataset from a visual-scene and auditory-linguistic N400 study. Zenodo. doi:10.5281/zenodo.1421347.

Draschkow, D., Heikel, E., Võ, M.L.H., Fiebach, C.J., Sassenhagen, J., 2018b. No evidence from MVPA for different processes underlying the N300 and N400 incongruity effects in object-scene processing. Neuropsychologia 120, 9–17. doi:10.1016/j.neuropsychologia.2018.09.016.

Gramfort, A., Luessi, M., Larson, E., Engemann, D.A., Strohmeier, D., Brodbeck, C., Goj, R., Jas, M., Brooks, T., Parkkonen, L., Hämäläinen, M., 2013. MEG and EEG data analysis with MNE-Python. Frontiers in Neuroscience 7, 267. doi:10.3389/fnins.2013.00267.

Grootswagers, T., Wardle, S.G., Carlson, T.A., 2017. Decoding dynamic brain patterns from evoked responses: A tutorial on multivariate pattern analysis applied to time series neuroimaging data. Journal of Cognitive Neuroscience 29, 677–697. doi:10.1162/jocn_a_01068.

Guggenmos, M., Sterzer, P., Cichy, R.M., 2018. Multivariate pattern analysis for MEG: A comparison of dissimilarity measures. NeuroImage 173, 434–447. doi:10.1016/j.neuroimage.2018.02.044.

Hald, L.A., Bastiaansen, M.C.M., Hagoort, P., 2006. EEG theta and gamma responses to semantic violations in online sentence processing. Brain and Language 96, 90–105. doi:10.1016/j.bandl.2005.06.007.

He, Y., Sommer, J., Hansen-Schirra, S., Nagels, A., 2024. Multivariate pattern analysis of EEG reveals nuanced impact of negation on sentence processing in the N400 and later time windows. Psychophysiology 61, e14491. doi:10.1111/psyp.14491.

Heikel, E., Sassenhagen, J., Fiebach, C.J., 2018. Time-generalized multivariate analysis of EEG responses reveals a cascading architecture of semantic mismatch processing. Brain and Language 184, 43–53. doi:10.1016/j.bandl.2018.06.007.

Hollenstein, N., Rotsztejn, J., Troendle, M., Pedroni, A., Zhang, C., Langer, N., 2018. ZuCo, a simultaneous EEG and eye-tracking resource for natural sentence reading. Scientific Data 5, 180291. doi:10.1038/sdata.2018.291.

Holm, S., 1979. A simple sequentially rejective multiple test procedure. Scandinavian Journal of Statistics 6, 65–70.

Hou, L., Zhang, G., Hao, Y., Kärkkäinen, T., Cong, F., 2026. Task-dependent effects of channel density on multivariate pattern analysis and spatial representational similarity analysis: Evidence from three EEG datasets. bioRxiv doi:10.64898/2026.01.26.701731. preprint.

Jas, M., Engemann, D.A., Bekhti, Y., Raimondo, F., Gramfort, A., 2017. Autoreject: Automated artifact rejection for MEG and EEG data. NeuroImage 159, 417–429. doi:10.1016/j.neuroimage.2017.06.030.

Jo, H., Yang, Y., Han, J., Duan, Y., Xiong, H., Lee, W.H., 2024. Are EEG-to-text models working? arXiv preprint arXiv:2405.06459. doi:10.48550/arXiv.2405.06459.

Kappenman, E.S., Farrens, J.L., Zhang, W., Stewart, A.X., Luck, S.J., 2021. ERP CORE: An open resource for human event-related potential research. NeuroImage 225, 117465. doi:10.1016/j.neuroimage.2020.117465.

Karimi-Rouzbahani, H., Shahmohammadi, M., Vahab, E., Setayeshi, S., Carlson, T., 2021. Temporal variabilities provide additional category-related information in object category decoding: A systematic comparison of informative EEG features. Neural Computation 33, 3027–3072. doi:10.1162/neco_a_01436.

Kessler, R., Enge, A., Skeide, M.A., 2025. How EEG preprocessing shapes decoding performance. Communications Biology 8, 1039. doi:10.1038/s42003-025-08464-3.

Krugliak, A., Draschkow, D., Võ, M.L.H., Clarke, A., 2024. Semantic object processing is modulated by prior scene context. Language, Cognition and Neuroscience 39, 962–971. doi:10.1080/23273798.2023.2279083.

Kutas, M., Federmeier, K.D., 2011. Thirty years and counting: Finding meaning in the N400 component of the event-related brain potential (ERP). Annual Review of Psychology 62, 621–647. doi:10.1146/annurev.psych.093008.131123.

Kutas, M., Hillyard, S.A., 1980. Reading senseless sentences: Brain potentials reflect semantic incongruity. Science 207, 203–205. doi:10.1126/science.7350657.

Larson, E., Gramfort, A., Engemann, D., Leppakangas, J., Brodbeck, C., Jas, M., et al., 2026. MNE-Python. Zenodo, version 1.13.2. doi:10.5281/zenodo.22712317.

Lawhern, V.J., Solon, A.J., Waytowich, N.R., Gordon, S.M., Hung, C.P., Lance, B.J., 2018. EEGNet: A compact convolutional neural network for EEG-based brain–computer interfaces. Journal of Neural Engineering 15, 056013. doi:10.1088/1741-2552/aace8c.

Ledoit, O., Wolf, M., 2004. A well-conditioned estimator for large-dimensional covariance matrices. Journal of Multivariate Analysis 88, 365–411. doi:10.1016/S0047-259X(03)00096-4.

Luck, S.J., 2020. ERP CORE (Compendium of Open Resources and Experiments). University of California, Davis; Open Science Framework. doi:10.18115/D5JW4R.

Olejnik, S., Algina, J., 2003. Generalized eta and omega squared statistics: Measures of effect size for some common research designs. Psychological Methods 8, 434–447. doi:10.1037/1082-989X.8.4.434.

Pedregosa, F., Varoquaux, G., Gramfort, A., Michel, V., Thirion, B., Grisel, O., Blondel, M., Prettenhofer, P., Weiss, R., Dubourg, V., Vanderplas, J., Passos, A., Cournapeau, D., Brucher, M., Perrot, M., Duchesnay, É., 2011. Scikit-learn: Machine learning in Python. Journal of Machine Learning Research 12, 2825–2830.

Petit, S., Badcock, N.A., Grootswagers, T., Woolgar, A., 2020. Unconstrained multivariate EEG decoding can help detect lexical-semantic processing in individual children. Scientific Reports 10, 10849. doi:10.1038/s41598-020-67407-6.

Qin, Y., Xu, Q., Kujala, T., Wang, X., Cong, F., 2026. Evaluating spatial normalization for SVM-based EEG decoding: A within- and between-subjects perspective. Biomedical Signal Processing and Control 116, 109535. doi:10.1016/j.bspc.2026.109535.

Ritchie, J.B., Lee Masson, H., Bracci, S., Op de Beeck, H.P., 2021. The unreliable influence of multivariate noise normalization on the reliability of neural dissimilarity. NeuroImage 245, 118686. doi:10.1016/j.neuroimage.2021.118686.

Schirrmeister, R.T., Springenberg, J.T., Fiederer, L.D.J., Glasstetter, M., Eggensperger, K., Tangermann, M., Hutter, F., Burgard, W., Ball, T., 2017. Deep learning with convolutional neural networks for EEG decoding and visualization. Human Brain Mapping 38, 5391–5420. doi:10.1002/hbm.23730.

Smith, S.M., Nichols, T.E., 2009. Threshold-free cluster enhancement: Addressing problems of smoothing, threshold dependence and localisation in cluster inference. NeuroImage 44, 83–98. doi:10.1016/j.neuroimage.2008.03.061.

Tanaka, H., Watanabe, H., Maki, H., Sakti, S., Nakamura, S., 2019. Electroencephalogram-based single-trial detection of language expectation violations in listening to speech. Frontiers in Computational Neuroscience 13, 15. doi:10.3389/fncom.2019.00015.

Trammel, T., Khodayari, N., Luck, S.J., Traxler, M.J., Swaab, T.Y., 2023. Decoding semantic relatedness and prediction from EEG: A classification method comparison. NeuroImage 277, 120268. doi:10.1016/j.neuroimage.2023.120268.

Wang, Z., Ji, H., 2022. Open vocabulary electroencephalography-to-text decoding and zero-shot sentiment classification, in: Proceedings of the AAAI Conference on Artificial Intelligence, pp. 5350–5358. doi:10.1609/aaai.v36i5.20472.

Wimmer, M., Veas, E.E., Müller-Putz, G.R., 2024. EEG-based incongruency decoding in AR with sLDA, SVM, and EEGNet. Current Directions in Biomedical Engineering 10, 21–24. doi:10.1515/cdbme-2024-1106.

Wobbrock, J.O., Findlater, L., Gergle, D., Higgins, J.J., 2011. The aligned rank transform for nonparametric factorial analyses using only ANOVA procedures, in: Proceedings of the SIGCHI Conference on Human Factors in Computing Systems, pp. 143–146. doi:10.1145/1978942.1978963.

Wolpert, D.H., 1992. Stacked generalization. Neural Networks 5, 241–259. doi:10.1016/S0893-6080(05)80023-1.

Zhang, G., Carrasco, C.D., Winsler, K., Bahle, B., Cong, F., Luck, S.J., 2024. Assessing the effectiveness of spatial PCA on SVM-based decoding of EEG data. NeuroImage 293, 120625. doi:10.1016/j.neuroimage.2024.120625.

Zhang, G., Luck, S.J., 2025. Assessing the impact of artifact correction and artifact rejection on the performance of SVM- and LDA-based decoding of EEG signals. NeuroImage 316, 121304. doi:10.1016/j.neuroimage.2025.121304.

Zhang, G., Wang, X., Winsler, K., Luck, S.J., 2026a. Evaluating the effects of regularization and cross-validation parameters on the performance of SVM-based decoding of EEG data. Cortex 201, 171–189. doi:10.1016/j.cortex.2026.05.005.

Zhang, G., Wang, X., Xin, Y., Cong, F., He, W., Luo, W., 2026b. The impact of downsampling on data quality, univariate measurement and multivariate pattern analysis in event-related potential research. NeuroImage 334, 121981. doi:10.1016/j.neuroimage.2026.121981.

