## Supplementary Material for "Feature processing for single-trial semantic decoding from EEG: a factorial comparison with pre-registered replication"

Section, table and figure numbers without the prefix S refer to the main text. Differences in balanced accuracy are given in percentage points (pp). All tables are generated from the analysis results by the script `scripts/paper_supplement.py` in the code repository.

#### S1 Data and preprocessing

Table S1 summarizes the preprocessing of the main analysis. In ERP CORE, 57 epochs in which autoreject flagged every channel were removed; 40 of them came from one participant (sub-029), in whom large transient artifacts on the reference electrode P10 affected all channels after re-referencing. No participant was excluded in any task of the main analysis. In the sentence task, epochs were removed by the global peak-to-peak threshold rather than repaired (Section 2.2), which removed 641 epochs; in addition, 56 flat epochs, all from one partly flat recording, were removed beforehand (Section S7). The total of 6401 recorded epochs, rather than 6400, arises because one sentence was presented twice to one participant and both presentations were kept. In nine participants exactly 129 of 160 epochs were kept, because autoreject selects the threshold from a discrete set of candidate values and these participants fell on the same quantile.

**Replication sample** Of the 288 PURSUE participants, three recordings without EEG data were excluded in advance. One recording used a different 32-channel montage that lacked ten of the pre-registered electrodes and could not be processed as pre-registered; it was excluded as a deviation from the pre-registration. Of the remaining 284 participants, one had fewer than 30 trials in one class after preprocessing (32 and 23) and was excluded as pre-registered, leaving 283 (103, 76 and 104 at the three sites). In one recording the first target occurred 0.48 s after the start of the recording, so that its envelope window ($-0.5$ s) could not be extracted; this trial was removed from both the waveform and the envelope epochs. Without the post-preprocessing exclusion ($n = 284$), R1 and R2 were virtually unchanged ($-3.6$ and $+15.0$ pp).

**Table S1:** Preprocessing summary (main analysis). Ocular and muscle independent components (ICs) removed and fraction of interpolated channel-epochs: mean (range). The last three columns count rejected epochs: without any good channel, above the global peak-to-peak threshold, and flat (sentence task only); –, step not applied. PURSUE includes the participant excluded after preprocessing.

| Dataset | $n$ | Trials kept | Per participant | Ocular ICs | Muscle ICs | Interpolated | No good channel | Threshold | Flat |
|---|---|---|---|---|---|---|---|---|---|
| Word pairs | 40 | 4743 / 4800 (98.8%) | 80–120 | 1.9 (1–3) | 7.1 (0–16) | 0.178 (0.083–0.445) | 57 | 0 | 0 |
| Scenes | 40 | 6075 / 6080 (99.9%) | 151–152 | 1.3 (0–3) | 13.3 (6–19) | 0.137 (0.040–0.415) | 5 | 0 | 0 |
| Sentences | 40 | 5704 / 6401 (89.1%) | 84–160 | 1.2 (0–2) | – | – | 0 | 641 | 56 |
| PURSUE | 284 | 33869 / 34078 (99.4%) | 55–121 | 2.2 (0–4) | 10.3 (0–18) | 0.141 (0.023–0.509) | 209 | 0 | 0 |

#### S2 Preprocessing adjustments after the pilot check and sensitivity analysis

**Pilot check** Before the full analysis of each task, the baseline configuration was decoded in three participants. For the scene and sentence tasks, the first pilot check gave accuracies close to chance (logistic regression: scenes 0.549, sentences 0.531; LDA 0.517 and 0.506; ERP CORE 0.671). Event timing and class labels had been verified against the authors' epochs. To locate the cause, the condition difference over the centro-parietal region was measured after each preprocessing step in one participant (Table S2). In the scene task the N400 effect was large and unaffected by preprocessing. In the sentence task it was weak from the start and was further reduced by the muscle independent component analysis (15 components removed) and by extensive interpolation (37% of channel-epochs).

**Table S2:** Condition difference (incongruent − congruent, µV) over the centro-parietal region after each preprocessing step, pilot participant sub-001 (*t*-values in parentheses where computed). ICA, independent component analysis.

| Processed up to | Scenes, 300–400 / 400–500 ms | Sentences, 300–400 / 400–500 ms |
|---|---|---|
| Filter and reference | −4.42 (−2.9) / −6.57 (−4.4) | −3.71 (−1.8) / −3.32 (−1.5) |
| + ocular ICA | −4.28 / −6.51 | −3.43 (−2.0) / −3.74 (−2.1) |
| + muscle ICA | −5.00 / −7.12 (12 components) | −1.50 (−1.1) / −1.75 (−1.2) (15 components) |
| + autoreject interpolation | −4.90 / −6.92 (interpolated 0.14) | −0.75 (−0.6) / −0.66 (−0.5) (interpolated 0.37) |

**Adjustments** Three settings were changed, each once and based on this diagnosis; alternative settings were not compared on the data. (1) In both tasks, the saved epochs and all features were restricted to the 28 electrodes used for ERP CORE (independent component analysis and interpolation still used all channels), so that the channel set C1 was identical across tasks. (2) In the sentence task, the muscle independent component analysis was omitted. (3) In the sentence task, epochs exceeding a global peak-to-peak threshold were rejected instead of interpolated. The adjustments restored the N400 effect of the sentence task in the pilot participant (about −3 µV between 300 and 500 ms) but did not raise decoding accuracy in the repeated pilot check (scenes 0.552, sentences 0.522); both tasks were then decoded in all 40 participants, where accuracy was clearly above chance (Table 4). In hindsight, the pilot participant was not representative of the scene task, whose group-average N400 effect was only −1.0 µV.

**Sensitivity analysis** The full analysis of both tasks was re-run with the preprocessing specified before the pilot check: all 60 EEG channels (so that C1 comprised 60 channels in these tasks), muscle independent component analysis and autoreject interpolation in both tasks. The interpretation of the possible outcomes was fixed before the analysis was run. Accuracy at baseline did not change significantly (Table S3). For logistic regression, all eleven joint categories were unchanged; for LDA, only the category of P2 changed (Table S4). Under the original path, the partly flat sentence recording kept 67 epochs (37 and 30), because 93 epochs had every channel flagged.

#### S3 Analyses within each dataset

Tables S5 and S6 give the planned contrasts within each dataset, Table S7 the four-way analyses of variance, and Table S8 the accuracy of each representation in each time window. For P2, the best

**Table S3:** Baseline accuracy with the main and the originally specified preprocessing, and correlation of the configuration means between the two.

| Task | Model | Baseline | Main | Original | Difference (pp) | $p$ | $r$ (configurations) |
|---|---|---|---|---|---|---|---|
| Scenes | LR | `B1-W1-N2-C1` | 0.553 | 0.551 | −0.23 | 0.510 | 0.993 |
| Scenes | LDA | `B1-W1-N1-C1` | 0.551 | 0.545 | −0.58 | 0.142 | 0.992 |
| Sentences | LR | `B1-W1-N2-C1` | 0.570 | 0.561 | −0.91 | 0.118 | 0.914 |
| Sentences | LDA | `B1-W1-N1-C1` | 0.556 | 0.552 | −0.48 | 0.454 | 0.869 |

**Table S4:** Joint test with the main and the originally specified preprocessing for the scene and sentence tasks (ERP CORE unchanged). Categories as in Table 3 of the main text; the arrow marks the one changed category.

| Model | Contrast | Main (pp) | $p_{\text{Holm}}$ | Category | Original (pp) | $p_{\text{Holm}}$ | Category |
|---|---|---|---|---|---|---|---|
| LR | P1 | −1.75 | < 0.0001 | G | −1.67 | < 0.0001 | G |
| LR | P2 | +0.72 | 0.016 | G | +0.86 | 0.005 | G |
| LR | P3 | +5.93 | < 0.0001 | C | +5.79 | < 0.0001 | C |
| LR | P4 | +0.26 | 0.457 | N | +0.27 | 0.372 | N |
| LR | P5 | +0.02 | 1.000 | N | +0.02 | 1.000 | N |
| LR | P6 | −0.35 | 0.015 | N | −0.26 | 0.095 | N |
| LR | P7 | −0.34 | 0.004 | G | −0.32 | 0.005 | G |
| LR | P8 | +0.45 | 0.085 | N | +0.32 | 0.338 | N |
| LR | P9 | +0.08 | 1.000 | N | +0.25 | 1.000 | N |
| LR | P10 | +0.16 | 1.000 | N | −0.08 | 1.000 | N |
| LR | P11 | 0.00 | 1.000 | N | −0.11 | 1.000 | N |
| LDA | P1 | −2.58 | < 0.0001 | G | −2.53 | < 0.0001 | G |
| LDA | P2 | +0.67 | 0.108 | N | +0.92 | 0.003 | G ← |
| LDA | P3 | +6.04 | < 0.0001 | C | +5.85 | < 0.0001 | C |
| LDA | P4 | +0.20 | 1.000 | N | +0.19 | 1.000 | N |
| LDA | P5 | +0.02 | 1.000 | N | +0.01 | 1.000 | N |
| LDA | P6 | +0.03 | 1.000 | N | +0.11 | 1.000 | N |
| LDA | P7 | −0.14 | 0.120 | N | −0.13 | 0.173 | N |
| LDA | P8 | +0.35 | 0.612 | N | +0.19 | 1.000 | N |
| LDA | P9 | −0.01 | 1.000 | N | +0.26 | 1.000 | N |
| LDA | P10 | +0.14 | 1.000 | N | −0.15 | 1.000 | N |
| LDA | P11 | +0.10 | 1.000 | N | +0.17 | 1.000 | N |

single band chosen by the leave-one-participant-out rule was theta (B2) for all 40 participants in the word-pair task, theta (B2) for 25 and alpha (B3) for 15 in the scene task, and alpha (B3) for all 40 participants in the sentence task.

**Table S5:** Planned contrasts within each dataset, logistic regression; Holm correction across the eleven contrasts of a dataset.

| Task | Contrast | Difference, pp (95% CI) | $d_z$ | $p_{\text{Holm}}$ | $p_{\text{Holm}}$ (Wilcoxon) |
|---|---|---|---|---|---|
| Word pairs | P1 | −2.25 (−2.97, −1.53) | −1.00 | < 0.0001 | < 0.0001 |
| | P2 | +0.92 (−0.04, +1.89) | 0.31 | 0.358 | 0.422 |
| | P3 | +9.80 (+7.76, +11.84) | 1.54 | < 0.0001 | < 0.0001 |
| | P4 | +0.87 (+0.17, +1.58) | 0.39 | 0.118 | 0.295 |
| | P5 | −0.10 (−0.39, +0.20) | −0.11 | 1.000 | 1.000 |
| | P6 | −0.76 (−1.17, −0.35) | −0.59 | 0.005 | 0.010 |
| | P7 | −0.45 (−0.79, −0.11) | −0.42 | 0.093 | 0.285 |
| | P8 | +0.60 (−0.09, +1.29) | 0.28 | 0.437 | 0.472 |
| | P9 | +0.53 (−0.50, +1.56) | 0.16 | 1.000 | 1.000 |
| | P10 | +0.45 (−0.47, +1.37) | 0.16 | 1.000 | 1.000 |
| | P11 | −0.19 (−0.87, +0.50) | −0.09 | 1.000 | 1.000 |
| Scenes | P1 | −1.37 (−2.02, −0.72) | −0.68 | 0.001 | 0.001 |
| | P2 | +1.13 (+0.44, +1.81) | 0.53 | 0.017 | 0.011 |
| | P3 | +3.19 (+1.94, +4.43) | 0.82 | < 0.0001 | < 0.0001 |
| | P4 | −0.27 (−0.71, +0.17) | −0.20 | 1.000 | 1.000 |
| | P5 | +0.05 (−0.19, +0.28) | 0.06 | 1.000 | 1.000 |
| | P6 | +0.01 (−0.31, +0.34) | 0.01 | 1.000 | 1.000 |
| | P7 | −0.16 (−0.46, +0.14) | −0.17 | 1.000 | 1.000 |
| | P8 | +0.46 (−0.07, +0.99) | 0.28 | 0.588 | 0.266 |
| | P9 | −0.05 (−0.77, +0.67) | −0.02 | 1.000 | 1.000 |
| | P10 | −0.70 (−1.46, +0.06) | −0.30 | 0.552 | 0.741 |
| | P11 | +0.07 (−0.54, +0.68) | 0.04 | 1.000 | 1.000 |
| Sentences | P1 | −1.63 (−2.32, −0.95) | −0.76 | 0.0002 | 0.0001 |
| | P2 | +0.12 (−0.71, +0.94) | 0.04 | 1.000 | 1.000 |
| | P3 | +4.81 (+3.44, +6.18) | 1.13 | < 0.0001 | < 0.0001 |
| | P4 | +0.18 (−0.28, +0.63) | 0.13 | 1.000 | 1.000 |
| | P5 | +0.11 (−0.11, +0.34) | 0.16 | 1.000 | 1.000 |
| | P6 | −0.31 (−0.68, +0.06) | −0.27 | 0.671 | 0.718 |
| | P7 | −0.42 (−0.75, −0.09) | −0.41 | 0.124 | 0.187 |
| | P8 | +0.28 (−0.35, +0.90) | 0.14 | 1.000 | 1.000 |
| | P9 | −0.23 (−1.05, +0.59) | −0.09 | 1.000 | 1.000 |
| | P10 | +0.74 (−0.05, +1.54) | 0.30 | 0.516 | 0.654 |
| | P11 | +0.12 (−0.40, +0.65) | 0.08 | 1.000 | 1.000 |

#### S4 Joint test with shrinkage LDA

Because LDA is invariant to per-feature affine transformations, N1 replaces N2 in P6 and P7. With LDA, P1 was general and P3 consistent in direction, as with logistic regression; P2 and P7 were not significant (Table S9). Across tasks, the difference between LDA and logistic regression among the waveform configurations decreased with the number of features (correlation with its logarithm: $r = -0.42$, $-0.48$ and $-0.37$ for word pairs, scenes and sentences): LDA was slightly better with at most 50 features and worse with more than 500.

**Table S6:** Planned contrasts within each dataset, shrinkage LDA (N1 in place of N2); Holm correction across the eleven contrasts of a dataset.

| Task | Contrast | Difference, pp (95% CI) | $d_z$ | $p_{\mathrm{Holm}}$ | $p_{\mathrm{Holm}}$ (Wilcoxon) |
|---|---|---|---|---|---|
| Word pairs | P1 | −3.26 (−4.28, −2.23) | −1.02 | < 0.0001 | < 0.0001 |
| | P2 | +0.41 (−0.62, +1.44) | 0.13 | 1.000 | 1.000 |
| | P3 | +9.71 (+7.52, +11.90) | 1.42 | < 0.0001 | < 0.0001 |
| | P4 | +0.83 (+0.11, +1.55) | 0.37 | 0.220 | 0.542 |
| | P5 | −0.19 (−0.51, +0.14) | −0.18 | 1.000 | 1.000 |
| | P6 | −0.17 (−0.58, +0.25) | −0.13 | 1.000 | 1.000 |
| | P7 | −0.19 (−0.42, +0.05) | −0.25 | 0.921 | 1.000 |
| | P8 | +0.30 (−0.49, +1.09) | 0.12 | 1.000 | 1.000 |
| | P9 | +0.53 (−0.61, +1.67) | 0.15 | 1.000 | 1.000 |
| | P10 | +0.46 (−0.55, +1.47) | 0.15 | 1.000 | 1.000 |
| | P11 | −0.26 (−1.05, +0.54) | −0.10 | 1.000 | 1.000 |
| Scenes | P1 | −2.33 (−3.26, −1.41) | −0.81 | < 0.0001 | < 0.0001 |
| | P2 | +0.82 (−0.05, +1.70) | 0.30 | 0.509 | 0.681 |
| | P3 | +3.68 (+2.38, +4.98) | 0.91 | < 0.0001 | < 0.0001 |
| | P4 | −0.46 (−0.95, +0.03) | −0.30 | 0.509 | 0.542 |
| | P5 | +0.05 (−0.21, +0.31) | 0.06 | 1.000 | 1.000 |
| | P6 | +0.20 (−0.11, +0.50) | 0.21 | 1.000 | 0.681 |
| | P7 | −0.05 (−0.21, +0.10) | −0.11 | 1.000 | 1.000 |
| | P8 | +0.40 (−0.27, +1.08) | 0.19 | 1.000 | 0.681 |
| | P9 | −0.05 (−0.85, +0.76) | −0.02 | 1.000 | 1.000 |
| | P10 | −0.84 (−1.71, +0.02) | −0.31 | 0.496 | 0.542 |
| | P11 | +0.43 (−0.36, +1.21) | 0.17 | 1.000 | 1.000 |
| Sentences | P1 | −2.14 (−3.16, −1.12) | −0.67 | 0.001 | 0.0009 |
| | P2 | +0.77 (−0.14, +1.69) | 0.27 | 0.666 | 0.800 |
| | P3 | +4.74 (+3.34, +6.14) | 1.08 | < 0.0001 | < 0.0001 |
| | P4 | +0.23 (−0.28, +0.74) | 0.14 | 1.000 | 1.000 |
| | P5 | +0.18 (−0.10, +0.46) | 0.20 | 1.000 | 1.000 |
| | P6 | +0.07 (−0.28, +0.41) | 0.06 | 1.000 | 1.000 |
| | P7 | −0.17 (−0.35, +0.01) | −0.30 | 0.591 | 0.971 |
| | P8 | +0.36 (−0.37, +1.09) | 0.16 | 1.000 | 1.000 |
| | P9 | −0.51 (−1.54, +0.52) | −0.16 | 1.000 | 0.971 |
| | P10 | +0.81 (−0.09, +1.71) | 0.29 | 0.605 | 0.730 |
| | P11 | +0.12 (−0.54, +0.77) | 0.06 | 1.000 | 1.000 |

**Table S7:** Four-way repeated-measures analysis of variance within each dataset (logistic regression; Greenhouse–Geisser corrected; Benjamini–Hochberg correction over the four main effects and six two-way interactions) and the aligned-rank-transform check.

| Effect | Word pairs | | | | Scenes | | | | Sentences | | | |
|---|---|---|---|---|---|---|---|---|---|---|---|---|
| | $F$ | $p_{\mathrm{BH}}$ | $\eta^2_G$ | ART $p_{\mathrm{BH}}$ | $F$ | $p_{\mathrm{BH}}$ | $\eta^2_G$ | ART $p_{\mathrm{BH}}$ | $F$ | $p_{\mathrm{BH}}$ | $\eta^2_G$ | ART $p_{\mathrm{BH}}$ |
| B | 68.3 | < 0.0001 | 0.289 | < 0.0001 | 14.9 | < 0.0001 | 0.079 | < 0.0001 | 28.1 | < 0.0001 | 0.127 | < 0.0001 |
| W | 22.2 | < 0.0001 | 0.027 | < 0.0001 | 2.7 | 0.135 | 0.003 | 0.126 | 0.6 | 0.568 | 0.001 | 0.454 |
| N | 11.2 | < 0.0001 | 0.004 | < 0.0001 | 3.7 | 0.047 | 0.002 | 0.126 | 4.0 | 0.033 | 0.002 | 0.063 |
| C | 3.0 | 0.057 | 0.006 | 0.072 | 2.2 | 0.135 | 0.004 | 0.126 | 3.6 | 0.033 | 0.008 | 0.043 |
| B × W | 17.9 | < 0.0001 | 0.035 | < 0.0001 | 4.0 | 0.0008 | 0.011 | 0.008 | 1.0 | 0.499 | 0.002 | 0.454 |
| B × N | 8.7 | < 0.0001 | 0.007 | < 0.0001 | 4.0 | 0.0005 | 0.004 | 0.0007 | 3.7 | 0.002 | 0.004 | 0.002 |
| B × C | 2.0 | 0.030 | 0.005 | 0.072 | 2.0 | 0.047 | 0.008 | 0.081 | 2.1 | 0.033 | 0.007 | 0.063 |
| W × N | 5.7 | 0.0001 | 0.001 | 0.0002 | 1.3 | 0.271 | 0.000 | 0.168 | 3.7 | 0.007 | 0.001 | 0.002 |
| W × C | 0.7 | 0.647 | 0.000 | 0.517 | 1.6 | 0.200 | 0.002 | 0.126 | 0.8 | 0.568 | 0.001 | 0.596 |
| N × C | 3.4 | 0.007 | 0.001 | 0.008 | 0.6 | 0.706 | 0.000 | 0.791 | 2.1 | 0.071 | 0.001 | 0.067 |

**Table S8:** Mean balanced accuracy of each representation in each time window (logistic regression; each participant's accuracy averaged over normalization and channel set). W1, 200–800 ms; W2, 300–500 ms; W3, 500–800 ms; W4, 0–800 ms. B1, waveform; B2–B5, theta, alpha, beta and low-gamma envelopes; B6, B2–B5 concatenated; B7, relative band amplitudes; B8, B1 and B2–B5 concatenated.

| Representation | Word pairs | | | | Scenes | | | | Sentences | | | |
|---|---|---|---|---|---|---|---|---|---|---|---|---|
| | W1 | W2 | W3 | W4 | W1 | W2 | W3 | W4 | W1 | W2 | W3 | W4 |
| B1 | 0.639 | 0.625 | 0.560 | 0.637 | 0.540 | 0.532 | 0.520 | 0.542 | 0.552 | 0.544 | 0.545 | 0.552 |
| B2 | 0.520 | 0.517 | 0.510 | 0.523 | 0.502 | 0.502 | 0.501 | 0.501 | 0.501 | 0.501 | 0.498 | 0.501 |
| B3 | 0.512 | 0.502 | 0.507 | 0.515 | 0.500 | 0.498 | 0.508 | 0.499 | 0.502 | 0.503 | 0.502 | 0.504 |
| B4 | 0.520 | 0.515 | 0.506 | 0.515 | 0.498 | 0.503 | 0.499 | 0.499 | 0.502 | 0.502 | 0.499 | 0.498 |
| B5 | 0.515 | 0.512 | 0.514 | 0.513 | 0.493 | 0.509 | 0.487 | 0.497 | 0.494 | 0.496 | 0.500 | 0.499 |
| B6 | 0.535 | 0.521 | 0.519 | 0.532 | 0.498 | 0.503 | 0.495 | 0.497 | 0.503 | 0.502 | 0.504 | 0.507 |
| B7 | 0.528 | 0.517 | 0.514 | 0.527 | 0.494 | 0.501 | 0.495 | 0.494 | 0.504 | 0.502 | 0.504 | 0.505 |
| B8 | 0.611 | 0.601 | 0.549 | 0.610 | 0.523 | 0.522 | 0.510 | 0.523 | 0.535 | 0.528 | 0.528 | 0.537 |

**Table S9:** Joint test across the three tasks with shrinkage LDA (N1 in place of N2 in P6 and P7). Categories as in Table 3 of the main text.

| Contrast | Joint, pp (95% CI) | $d_z$ | $p_{\text{Holm}}$ | Word pairs | Scenes | Sentences | $p_{\text{het}}$ | Category |
|---|---|---|---|---|---|---|---|---|
| P1 | −2.58 (−3.12, −2.03) | −0.83 | < 0.0001 | −3.26 | −2.33 | −2.14 | 1.000 | G |
| P2 | +0.67 (+0.15, +1.19) | 0.23 | 0.108 | +0.41 | +0.82 | +0.77 | 1.000 | N |
| P3 | +6.04 (+5.09, +7.00) | 1.15 | < 0.0001 | +9.71 | +3.68 | +4.74 | 0.0001 | C |
| P4 | +0.20 (−0.12, +0.52) | 0.11 | 1.000 | +0.83 | −0.46 | +0.23 | 0.083 | N |
| P5 | +0.02 (−0.14, +0.17) | 0.02 | 1.000 | −0.19 | +0.05 | +0.18 | 1.000 | N |
| P6 | +0.03 (−0.17, +0.24) | 0.03 | 1.000 | −0.17 | +0.20 | +0.07 | 1.000 | N |
| P7 | −0.14 (−0.25, −0.03) | −0.23 | 0.120 | −0.19 | −0.05 | −0.17 | 1.000 | N |
| P8 | +0.35 (−0.05, +0.76) | 0.15 | 0.612 | +0.30 | +0.40 | +0.36 | 1.000 | N |
| P9 | −0.01 (−0.60, +0.58) | 0.00 | 1.000 | +0.53 | −0.05 | −0.51 | 1.000 | N |
| P10 | +0.14 (−0.37, +0.65) | 0.05 | 1.000 | +0.46 | −0.84 | +0.81 | 0.226 | N |
| P11 | +0.10 (−0.32, +0.51) | 0.04 | 1.000 | −0.26 | +0.43 | +0.12 | 1.000 | N |

#### S5 EEGNet

EEGNet (Lawhern et al., 2018) was implemented in braindecode (Aristimunha et al., 2026; Schirrmeister et al., 2017) and trained per participant with 5-fold cross-validation and three random seeds. Its input was the epoch restricted to the time window and channel set of the configuration, without averaging in 50-ms bins, so that only configurations with a time-series representation (B1–B5) could be evaluated. For each task, the baseline and the ten best and three worst of these configurations for logistic regression were run. EEGNet did not exceed logistic regression on any waveform configuration (Table S10). At baseline, logistic regression minus EEGNet was +1.5 pp ($p = 0.07$), +0.3 pp ($p = 0.64$) and +1.8 pp ($p = 0.036$) for word pairs, scenes and sentences; participant-level accuracies of the two models correlated with $r = 0.85$, 0.70 and 0.54, and the rank correlations of the 13 configuration means were 0.64, 0.88 and 0.63.

#### S6 Regularization grid

The regularization parameter of logistic regression often reached an end of the study grid. All 640 configurations of the three tasks were therefore re-run on the same folds with a grid extended by two orders of magnitude at each end. The extended grid did not improve accuracy (Table S11), accuracy was flat on the strongly regularized side, and the significance of all 33 within-dataset planned contrasts was unchanged. In the joint test, ten of the eleven categories were unchanged; the joint effect of P6 was significant with both grids ($p_{\text{Holm}} = 0.015$ and 0.008), and its category changed from no reliable general effect to general because its near-zero estimate in the scene task changed sign (+0.01 to −0.04 pp).

#### S7 The partly flat recording and the combination of bands (P2)

In the sentence recording of one participant (sub-017), all channels had abnormally low amplitude in about 25–30% of the recording (channel standard deviation about 1 µV, compared with about 10 µV normally), and about 10% contained large artifacts. The global peak-to-peak threshold was therefore estimated at 2.8 µV, so that only 33 flat epochs were kept and all normal epochs were rejected. This was corrected after the full analysis by removing flat epochs (median channel standard deviation below 1 µV) before the threshold was estimated; the participant then kept 84 epochs (39 and 45), and the threshold and kept epochs of all other recordings were unchanged. Before the correction, P2 was general for both classifiers (+1.2 pp for logistic regression); after it, only for logistic regression (+0.7 pp). Without this participant, P2 was no longer significant, whereas P1, P3 and P7 were unaffected (Table S12). P2 is also structurally fragile: if two bands have nearly equal marginal means, as alpha and theta did in the sentence task, removing a single participant can change the band chosen for all remaining participants. P2 is therefore not taken as a conclusion.

#### S8 Exploratory analyses

These analyses were carried out after the results were known. Table S13 gives the condition differences over the centro-parietal region. In ERP CORE, all band-envelope differences were within ±0.016 in $\log_{10}$ amplitude (less than 4% in amplitude), and none survived correction for the eight comparisons; the late increase in theta amplitude had the direction reported for semantic violations (Hald et al., 2006), and beta amplitude decreased in the same window. Table S14 gives correlations across

**Table S10:** EEGNet compared with logistic regression on the baseline and the ten best and three worst time-series configurations of each task (mean balanced accuracy).

| Task | Configuration | EEGNet | Logistic regression | Difference (pp) |
|---|---|---|---|---|
| Word pairs | `B1-W4-N3-C2` | 0.655 | 0.657 | −0.10 |
| | `B1-W4-N1-C2` | 0.652 | 0.656 | −0.39 |
| | `B1-W4-N2-C1` | 0.649 | 0.658 | −0.90 |
| | `B1-W1-N2-C1` | 0.648 | 0.663 | −1.45 |
| | `B1-W4-N3-C5` | 0.646 | 0.653 | −0.71 |
| | `B1-W4-N2-C5` | 0.646 | 0.655 | −0.97 |
| | `B1-W1-N2-C5` | 0.644 | 0.658 | −1.39 |
| | `B1-W4-N3-C1` | 0.642 | 0.655 | −1.26 |
| | `B1-W1-N3-C1` | 0.639 | 0.658 | −1.85 |
| | `B1-W2-N2-C1` | 0.619 | 0.654 | −3.47 |
| | `B4-W3-N4-C4` | 0.498 | 0.492 | +0.62 |
| | `B3-W2-N3-C5` | 0.489 | 0.489 | +0.08 |
| | `B3-W2-N3-C2` | 0.483 | 0.488 | −0.44 |
| Scenes | `B1-W1-N2-C1` | 0.550 | 0.553 | −0.27 |
| | `B1-W1-N3-C1` | 0.547 | 0.554 | −0.70 |
| | `B1-W4-N2-C1` | 0.542 | 0.553 | −1.12 |
| | `B1-W4-N2-C5` | 0.541 | 0.547 | −0.57 |
| | `B1-W4-N1-C1` | 0.539 | 0.551 | −1.21 |
| | `B1-W4-N1-C5` | 0.539 | 0.546 | −0.71 |
| | `B1-W4-N3-C1` | 0.539 | 0.553 | −1.38 |
| | `B1-W4-N4-C5` | 0.537 | 0.548 | −1.07 |
| | `B1-W4-N4-C3` | 0.533 | 0.547 | −1.42 |
| | `B1-W1-N4-C3` | 0.532 | 0.547 | −1.47 |
| | `B5-W3-N2-C3` | 0.483 | 0.479 | +0.42 |
| | `B5-W3-N4-C5` | 0.479 | 0.478 | +0.15 |
| | `B5-W3-N3-C5` | 0.479 | 0.479 | +0.01 |
| Sentences | `B1-W1-N2-C5` | 0.561 | 0.566 | −0.45 |
| | `B1-W1-N3-C5` | 0.558 | 0.563 | −0.45 |
| | `B1-W4-N3-C1` | 0.557 | 0.566 | −0.95 |
| | `B1-W4-N3-C5` | 0.557 | 0.562 | −0.54 |
| | `B1-W1-N3-C1` | 0.554 | 0.566 | −1.23 |
| | `B1-W4-N2-C1` | 0.553 | 0.564 | −1.16 |
| | `B1-W1-N2-C1` | 0.552 | 0.570 | −1.80 |
| | `B1-W1-N4-C5` | 0.551 | 0.565 | −1.36 |
| | `B1-W2-N2-C1` | 0.550 | 0.563 | −1.32 |
| | `B1-W4-N4-C5` | 0.547 | 0.562 | −1.51 |
| | `B5-W2-N4-C3` | 0.491 | 0.485 | +0.60 |
| | `B5-W2-N3-C3` | 0.489 | 0.481 | +0.82 |
| | `B5-W1-N4-C4` | 0.485 | 0.485 | +0.02 |

**Table S11:** Logistic regression with the study grid ($10^{-3}$ to $10^{2}$) and the extended grid ($10^{-5}$ to $10^{4}$) on the same folds: mean balanced accuracy at baseline and over all 640 configurations, and the correlation $r$ of the 640 configuration means between the two grids. The last two columns give the fraction of outer folds in which the extended grid selected a value outside the study grid.

| | Baseline | | | All 640 configurations | | | Outside study grid | |
|---|---|---|---|---|---|---|---|---|
| Task | Study | Extended | Difference (pp) | Study | Extended | $r$ | $C < 10^{-3}$ | $C > 10^{2}$ |
| Word pairs | 0.663 | 0.658 | −0.52 | 0.539 | 0.538 | 0.999 | 20.5% | 11.2% |
| Scenes | 0.553 | 0.548 | −0.54 | 0.506 | 0.506 | 0.989 | 25.6% | 11.6% |
| Sentences | 0.570 | 0.568 | −0.24 | 0.511 | 0.511 | 0.987 | 25.3% | 11.1% |

**Table S12:** Joint test (logistic regression) without the partly flat sentence recording (Draschkow participant sub-017, removed from both of that participant's tasks) and, in addition, without ERP CORE sub-029 (40 epochs without a good channel) and sub-040 (excluded in the original ERP CORE analyses). Each cell: estimate (pp), $p_{\mathrm{Holm}}$, category.

| Sample | P1 | P2 | P3 | P7 |
|---|---|---|---|---|
| All participants | −1.75, < 0.0001, G | +0.72, 0.016, G | +5.93, < 0.0001, C | −0.34, 0.004, G |
| Without sub-017 (scene and sentence) | −1.74, < 0.0001, G | +0.42, 0.377, N | +5.93, < 0.0001, C | −0.35, 0.004, G |
| Also without ERP CORE sub-029, sub-040 | −1.84, < 0.0001, G | +0.42, 0.401, N | +6.17, < 0.0001, C | −0.36, 0.003, G |

participants: in ERP CORE, the individual N400 effect explained about half of the variance in baseline accuracy, which was not the case for the late effect.

**Table S13:** Exploratory condition differences (incongruent − congruent) over the centro-parietal region: waveform in microvolts, band envelopes in $\log_{10}$ amplitude (0.016 corresponds to about 4%). Uncorrected one-sample *t*-tests across participants. Envelope differences were computed for ERP CORE only.

| Measure | Word pairs | | Scenes | | Sentences | |
|---|---|---|---|---|---|---|
| | Effect | *p* | Effect | *p* | Effect | *p* |
| Waveform, 300–500 ms (µV) | −5.28 | < 0.0001 | −0.97 | 0.0005 | −2.95 | < 0.0001 |
| Waveform, 500–800 ms (µV) | −2.07 | < 0.0001 | +0.17 | 0.533 | −1.18 | 0.010 |
| Theta envelope, 300–500 ms ($\log_{10}$) | −0.010 | 0.188 | – | – | – | – |
| Alpha envelope, 300–500 ms | +0.014 | 0.065 | – | – | – | – |
| Beta envelope, 300–500 ms | −0.009 | 0.137 | – | – | – | – |
| Low-gamma envelope, 300–500 ms | −0.003 | 0.315 | – | – | – | – |
| Theta envelope, 500–800 ms | +0.015 | 0.044 | – | – | – | – |
| Alpha envelope, 500–800 ms | +0.001 | 0.827 | – | – | – | – |
| Beta envelope, 500–800 ms | −0.014 | 0.010 | – | – | – | – |
| Low-gamma envelope, 500–800 ms | −0.003 | 0.323 | – | – | – | – |

#### S9 Time–frequency sensitivity analysis

**Decomposition** Complex Morlet wavelets were applied to the continuous preprocessed data at 12 logarithmically spaced frequencies (2.00, 2.60, 3.38, 4.40, 5.72, 7.44, 9.68, 12.6, 16.4, 21.3, 27.7 and 36.0 Hz), with the number of cycles increasing logarithmically from 3 to 10 (temporal standard deviation from 239 ms at 2 Hz to 44 ms at 36 Hz). Segments separated by boundary annotations were transformed separately. Epochs from −0.5 s to the end of the waveform epoch were extracted and the flagged channels interpolated on the real and imaginary parts, as for the band envelopes. Amplitude features were $\log_{10}|Z|$ baseline-corrected between −500 and −200 ms; phase features were $\cos\varphi$ and $\sin\varphi$ without baseline correction; both were averaged in 50-ms bins.

**Stacking** Within each outer training fold, each base model (B1 and T1, logistic regression) was tuned as in the main analysis and refitted on the whole training fold. With its regularization parameter fixed to the selected value, out-of-sample decision values for the training trials were obtained by stratified 5-fold cross-fitting. These were standardized within the training fold and combined by a logistic-regression meta-model ($C = 1$). Test trials were classified by the meta-model from the decision values of the refitted base models. The regularization parameter was fixed during cross-fitting and the decision values standardized because, in a preliminary test on one participant before the main run, re-tuning the parameter in each cross-fitting fold gave decision values whose scale differed between cross-fitting and testing, so that the meta-model assigned most test trials

**Table S14:** Exploratory correlations across participants ($n = 40$ per task). N400 and late effect, incongruent − congruent over the centro-parietal region at 300–500 and 500–800 ms, in µV or as a single-trial effect size $d$; LR, logistic regression.

| Task | Variables | Pearson $r$ | $p$ | Spearman $\rho$ |
|---|---|---|---|---|
| Word pairs | N400 effect (µV) vs. baseline LR accuracy | −0.73 | < 0.0001 | −0.72 |
| | N400 effect ($d$) vs. baseline LR accuracy | −0.71 | < 0.0001 | −0.70 |
| | Late effect ($d$) vs. baseline LR accuracy | −0.13 | 0.411 | −0.10 |
| | B1 vs. mean single-band envelope accuracy (LR marginal means) | 0.47 | 0.002 | 0.54 |
| | EEGNet vs. LR accuracy at the baseline configuration | 0.85 | < 0.0001 | 0.84 |
| Scenes | N400 effect (µV) vs. baseline LR accuracy | −0.12 | 0.448 | −0.17 |
| | N400 effect ($d$) vs. baseline LR accuracy | −0.14 | 0.391 | −0.16 |
| | Late effect ($d$) vs. baseline LR accuracy | −0.09 | 0.583 | −0.13 |
| | B1 vs. mean single-band envelope accuracy (LR marginal means) | −0.42 | 0.007 | −0.43 |
| | EEGNet vs. LR accuracy at the baseline configuration | 0.70 | < 0.0001 | 0.73 |
| Sentences | N400 effect (µV) vs. baseline LR accuracy | −0.35 | 0.027 | −0.41 |
| | N400 effect ($d$) vs. baseline LR accuracy | −0.35 | 0.026 | −0.41 |
| | Late effect ($d$) vs. baseline LR accuracy | 0.00 | 0.977 | −0.03 |
| | B1 vs. mean single-band envelope accuracy (LR marginal means) | 0.01 | 0.945 | 0.03 |
| | EEGNet vs. LR accuracy at the baseline configuration | 0.54 | 0.0003 | 0.53 |

to one class. Table S15 gives the mean accuracy of each representation and the stacking weights. The comparisons F1–F6 of the main text are labelled S1–S6 in the pre-specification and the code repository; these labels are unrelated to the numbering of the supplementary sections.

**Table S15:** Time–frequency sensitivity analysis: mean balanced accuracy of each representation (logistic regression; other dimensions at baseline), the fraction of outer folds in which the stacking meta-model gave wavelet power (T1) a positive weight, and the median share of T1, $|w_{T1}|/(|w_{B1}| + |w_{T1}|)$.

| Task | B1 | B6 | T1 | T2 | T3 | T4 | T5 | $w_{T1} > 0$ | Share of T1 |
|---|---|---|---|---|---|---|---|---|---|
| Word pairs | 0.663 | 0.550 | 0.568 | 0.573 | 0.651 | 0.612 | 0.659 | 78% | 0.22 |
| Scenes | 0.553 | 0.498 | 0.514 | 0.511 | 0.524 | 0.526 | 0.529 | 58% | 0.13 |
| Sentences | 0.570 | 0.509 | 0.520 | 0.517 | 0.519 | 0.546 | 0.556 | 58% | 0.15 |

### References


Aristimunha, B., Guetschel, P., Wimpff, M., Gemein, L., Rommel, C., Banville, H., Sliwowski, M., Wilson, D., Brandt, S., Gnassounou, T., Paillard, J., Srivastava, A., Junqueira Lopes, B., Sedlar, S., Tayal, S., Truong, Y., Moreau, T., Chevallier, S., Gramfort, A., Schirrmeister, R.T., 2026. Braindecode: Toolbox for decoding raw electrophysiological brain data with deep learning models. Zenodo, version 1.8.1. doi:10.5281/zenodo.22209839.

Hald, L.A., Bastiaansen, M.C.M., Hagoort, P., 2006. EEG theta and gamma responses to semantic violations in online sentence processing. Brain and Language 96, 90–105. doi:10.1016/j.bandl.2005.06.007.

Lawhern, V.J., Solon, A.J., Waytowich, N.R., Gordon, S.M., Hung, C.P., Lance, B.J., 2018. EEGNet: A compact convolutional neural network for EEG-based brain–computer interfaces. Journal of Neural Engineering 15, 056013. doi:10.1088/1741-2552/aace8c.

Schirrmeister, R.T., Springenberg, J.T., Fiederer, L.D.J., Glasstetter, M., Eggensperger, K.,

Tangermann, M., Hutter, F., Burgard, W., Ball, T., 2017. Deep learning with convolutional neural networks for EEG decoding and visualization. Human Brain Mapping 38, 5391–5420. doi:10.1002/hbm.23730.
